# A sparse neural code for acoustic targets enables speech-like vocal flexibility in parrots

**DOI:** 10.64898/2026.10.21.760739

**Authors:** Zhilei Zhao, Alison N. Plump, Caleb Jones, Han Kheng Teoh, Brian Kardon, Jesse H. Goldberg

## Abstract

English speech flexibly recombines 44 phonemes into >20,000 words and limitless phrases. Here we identify a novel neural code supporting combinatorial vocal control in budgerigar parrots. Deep learning based acoustic analyses show that both English imitated and natural warble songs are composed of subsyllabic acoustic elements that are flexibly reused across acoustic contexts, analogous to phoneme reuse in words. Such reuse poses a scaling problem for songbird-inspired timing-based codes, which require distinct neural sequences for distinct syllables even when syllables share acoustic components. We identified neurons in the parrot frontal cortical nucleus MO that exhibit sparse bursts locked to reused acoustic elements independent of syllable identity, timing, or surrounding acoustic context. These results provide the first evidence for a sparse, acoustically-grounded vocal motor code that can be flexibly reused to assemble a large and combinatorial vocal repertoire.

## Main text

In humans, the vast vocal repertoire arises from the combinatorial power with which a finite set of reusable acoustic elements, or phonemes, can be flexibly sequenced^1^. Speech can be viewed as a trajectory through an acoustic space of phonemes^2,3^. For example, the words “bird” and “burst” share the same path until they diverge at the final consonant cluster (Fig. 1a). Neural mechanisms underlying flexible and seamless navigation through acoustic space are poorly understood.

**Figure 1.**
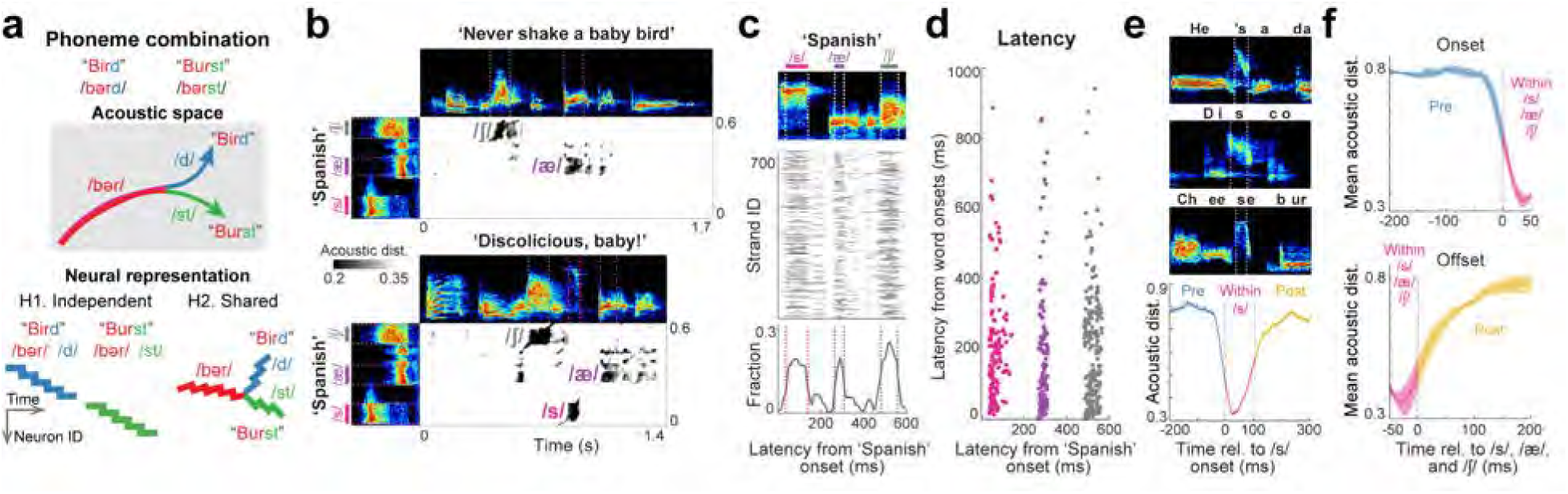
Flexible reuse of subsyllabic acoustic elements in an English-imitating budgerigar. a, Two hypotheses for neural control of vocal production. Words that share acoustic elements (for example “bird” and “burst”) could be generated by independent neural sequences despite shared acoustics (H1, songbird inspired), or could share neural representations where acoustics overlap (H2). b, Latent space distance metrics identify reused acoustic elements in an acoustic similarity matrix. Top, spectrograms of example English sentences produced by Disco. Left, spectrogram of Disco uttering ‘Spanish.’ Middle, acoustic similarity matrix between example sentences and the reference word. Black strands indicate intervals with high acoustic similarity. c, Unsupervised discovery of reused acoustic primitives. Top, spectrogram of the reference “Spanish.” Middle, high similarity strands obtained by scanning the reference against the full Disco dataset, plotted in reference time coordinates. Bottom, strand density across reference time. Peaks in density define candidate reused primitives (dashed lines). d, Variable timing of reused primitives. Latency of the three example primitives in the reference utterance (x axis) and in matched segments across the dataset (y axis). Each point denotes the initiation time from syllable onset of one high similarity strand. e, Convergence and divergence around a reused primitive. Top, example phrases containing a match to the primitive highlighted in c (pink). Bottom, mean pairwise acoustic distance aligned to primitive onset across all matches, showing transient convergence during the primitive and divergence before and after. f, Same analysis as e for all three primitives identified in c. Solid lines indicate mean and shading indicates SEM.

### Flexible reuse of subsyllabic acoustic elements in an English-imitating budgerigar

Budgerigar parrots (*Melopsittacus undulatus*) may have the largest vocal repertoire of any nonhuman animal and may hold a record for the most English-imitated words^4–8^. To test whether speech-like reuse of subsyllabic acoustic elements contributes to repertoire size, we analyzed the vocalizations of an English-imitating budgerigar named Disco. A human annotator and OpenAI’s Whisper network independently transcribed 26 minutes of vocalizations. Grapheme-to-phoneme (G2P) analysis of concordant annotations suggested that Disco reused 37 phonemes across 179 distinct words (Methods). However, because both human listeners and speech recognition systems are biased toward human language categories, this analysis alone cannot establish phoneme-like reuse on purely acoustic grounds.

In songbirds, gap-bounded syllables are generally considered the fundamental building blocks of vocal communication^7,9^. By contrast, Disco’s imitated English, like human speech, contained acoustic elements that were flexibly recombined within syllables. To test for reuse of subsyllabic acoustic elements without linguistic priors, we represented vocalizations as continuous trajectories through a low-dimensional acoustic space using a ‘shotgun’ variational autoencoder (VAE) trained on short, overlapping spectrogram windows slid through the Disco dataset (32 ms window, 1 ms step; Methods)^10^. This approach enabled comparison of fine-scale acoustic structure across vocalizations independent of syllable boundaries.

Our goal was not to construct a complete inventory of discrete acoustic elements, but rather to identify clear examples and test whether such elements could be flexibly reused across imitated words and syllables. To this end, we compared the acoustic trajectory from a single instance of Disco saying “Spanish” to trajectories from Disco’s full repertoire (n = 2931 syllables). Short stretches of sound were classified as reused matches when they exhibited high similarity for a minimum duration in the acoustic cross-correlation matrix (cosine distance < 0.2 for > 20 milliseconds; Methods). Using this approach, the /s/, /æ/, and /ʃ/ sounds in the reference utterance were detected in phrases such as “Never shake a baby bird” and “Discolicious baby” (Fig. 1b-c).

Across the entire dataset, latent trajectories corresponding to the /s/, /æ/, and /ʃ/ sounds in the reference “Spanish” utterance were matched 141, 148, and 224 times, respectively. Importantly, these reused elements occurred at variable latencies relative to syllable onset (Fig. 1d) and in diverse acoustic contexts. For example, reuses of the /s/-like element in “He’s a dabba dabba doo bird,” “Disco,” and “Cheeseburger” were detected without supervision (Fig. 1e). Pairwise acoustic distance measurements revealed a brief convergence in acoustic structure at the onset of each reused element, followed by rapid divergence thereafter (Fig. 1e-f), demonstrating that Disco’s vocalizations can approach and exit the same acoustic target along different paths. This trajectory-based reuse is a hallmark of human speech^2,3^. Together, these analyses show that representing sounds as trajectories in a VAE-derived acoustic space enables unsupervised identification and quantification of reused phoneme-like acoustic elements in nonhuman vocalizations. Similar unsupervised approaches have been used to discover phoneme-like units in human speech without labeled training data^11^.

### Flexible reuse of subsyllabic acoustic elements in natural budgerigar songs

To test if budgerigars also exhibit phoneme-like acoustic element reuse in their natural vocalizations, we conducted the same analyses on warble songs produced during face-to-face social interactions. Natural budgerigar warble consists of discrete and gap-bounded ‘elemental’ syllables (calls, clicks, squawks) as well as long, gapless ‘compound’ syllables proposed to share structural features with gapless speech^7,8,12^ (Fig. 2a). We fine-tuned a transformer-based segmentation network, WhisperSeg^13^ to label calls, clicks, squawks and compound syllables on a large corpus of warble songs; highly variable syllables uncategorizable by humans or the network were collectively labeled ‘X’ (Fig. 2a-b, Extended Data Fig. 1, Methods, n=5916 syllables from 4 birds for model training).

**Figure 2.**
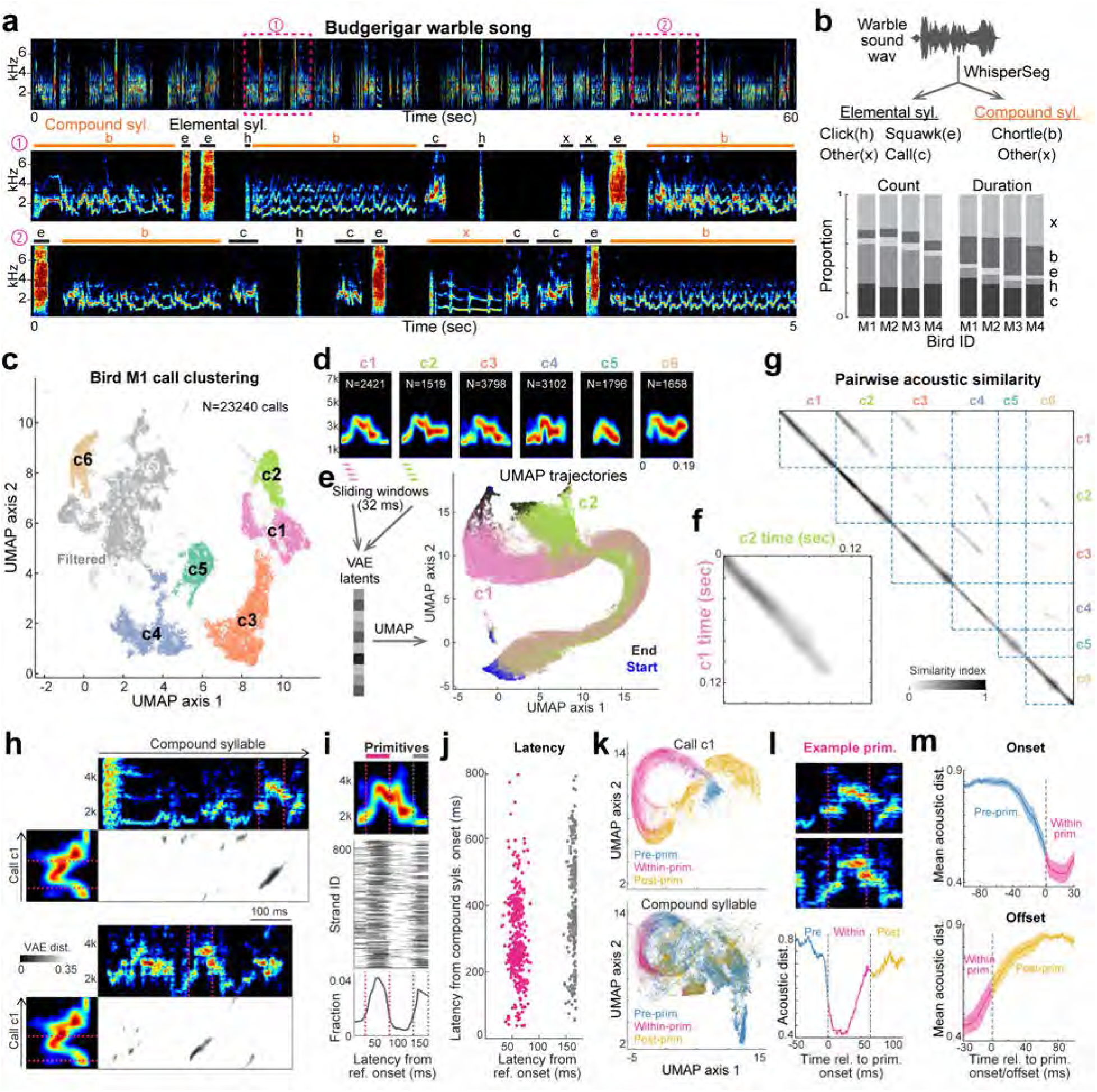
Flexible reuse of subsyllabic acoustic elements in natural budgerigar warble songs. a, Example warble song spectrogram. Top, a 60 s segment from a warble episode lasting more than 10 min. Middle and bottom, two 5 s zoomed-in segments. Letters indicate syllable types defined in b. b, Warble analysis pipeline and syllable composition. Top, schematic of syllable-level segmentation and categorization. Bottom, proportion of each syllable type by syllable count (left) and total duration (right). c, UMAP projection of calls produced during bird M1 warble. Each point represents one call, colored by HDBSCAN cluster label. Light grey denotes unclustered calls; dark grey denotes a call cluster excluded by quality control (Methods). d, Averaged spectrograms of identified call subtypes from bird M1, colored as in c. Numbers indicate the number of renditions contributing to each average. e, Left, schematics of the shotgun VAE analysis. Right, UMAP trajectories of 500 renditions of c1 and c2 calls. Blue dots indicate syllable onsets, black dots indicate syllable offsets. f, Acoustic similarity between all pairs of time points in call subtypes c1 and c2, quantified using the normalized dMMD metric (Methods). High similarity is observed during the initial and middle portions of the calls, with divergence toward the ends. g, Acoustic similarity matrices as in f for all pairwise comparisons between call subtypes in bird M1. h, Identification of shared vocal primitives between calls and compound syllables. Top, spectrograms of example compound syllables. Left, mean spectrogram of call subtype c1 (bird M1), used as the reference. Middle, acoustic distance matrix between the compound syllable and the reference call. Black strands indicate regions of high acoustic similarity. i, Localization of shared primitives within the reference call. Top, mean spectrogram of call subtype c1. Middle, high-similarity strands from call–compound comparisons, plotted in reference time coordinates. Bottom, strand density across reference time; peaks (dashed lines) define reused primitives. j, Variable timing of reused primitives. Latency of the two example primitives in the reference call (x axis) and in compound syllables (y axis). Each point represents one high-similarity strand. k, UMAP embedding of spectrogram windows surrounding the shared primitive (pink in panel i) in the reference call (top) and compound syllables (bottom). Each point corresponds to a 32 ms sliding window projected into acoustic space. l, Convergence and divergence of acoustic trajectories around a shared primitive. Primitive onset aligned spectrograms (top) and mean pairwise acoustic distance (bottom). m, Same analysis as l for all identified primitives. Solid lines denote mean; shaded regions indicate SEM.

If budgerigar vocalizations are combinatorial, then even simpler elemental calls may be composed of smaller subsyllabic acoustic elements, just as brief words in speech (e.g., burst, bird) arise from recombination of discrete phonemes^1,2^. Calls constituted roughly one third of all warble syllables (range: 23–32%, n=4 birds) and could be clustered into distinct types when VAEs were trained to reconstruct each call spectrogram as a single image (n=3-6 types, Fig. 2c-d, Extended Data Figs. 1-2, Methods), as shown previously^6,14^. Yet even in distinct call types visual inspection of spectrograms suggested that subsyllabic elements could be shared across distinct call types. For example, call variants c1 and c2 produced by bird M1 yielded distinguishable clusters in the UMAP projections of their VAE latents (Fig. 2c), but both began with nearly identical frequency contours but diverged at their termini: c1 ended in a soft downsweep to 1 kHz, whereas c2 rose sharply to 3 kHz (Fig. 2d). To visualize reuse of acoustic elements across call types, we examined UMAP projections of latent acoustic trajectories derived from a shotgun VAE trained on sliding spectrogram windows through all c1 and c2 renditions (32 ms window, 1 ms step, Methods, Fig. 2e). Like ‘bird’ and ‘burst,’ both call types followed a shared path through latent acoustic space before diverging toward distinct endpoints (Fig. 2e). Maximum mean discrepancy (MMD) analysis^10^ was used to compute the acoustic distance between call types and confirmed that the first ∼100 ms of c1 and c2 were acoustically similar but diverged sharply thereafter (Fig. 2f, Methods). We quantified reuse of subsyllabic acoustic trajectories by repeating pairwise comparisons across all call types in each bird and, as with our approach to Disco, conservatively defined a reused trajectory as strands with cosine distance < 0.2 and duration > 20 ms (Fig. 2g, Methods). In 30 pairwise comparisons across 4 birds, 53.3% of call pairs shared at least one subsyllabic acoustic trajectory, comprising 29.5% of total call duration. Thus, even calls, the simplest of learned budgerigar vocalizations^6^, exhibited a combinatorial structure based on reuse of smaller subsyllabic acoustic elements.

To test for acoustic element reuse in compound syllables, we used calls as acoustic templates and scanned their latent embeddings through all compound syllables (n=37743 syllables in 4 birds). In the acoustic similarity matrix, we identified high-similarity strands between template calls and individual compound syllables (cosine distance<0.2, duration>20 ms, Fig. 2h). When these strands were mapped back to the template call, they concentrated in certain parts of the call (Fig. 2i, Extended Data Fig. 3), which we termed “vocal primitives” to indicate high reuse frequency. Notably, like ‘phonemes’ in Disco’s imitated speech, primitives occurred with a highly variable latency relative to compound syllable onset (Fig. 2j). In addition, across renditions, a given primitive was preceded and followed by diverse sounds, as quantified by pairwise acoustic distance (Fig. 2k-m). Thus, just as there were many paths leading into or out of the /s/, /æ/, and /ʃ/ sounds in Disco’s imitated English, natural warble song contained acoustic primitives that could be entered or exited via many routes in acoustic space.

### Identifying a candidate ‘state space’ for vocal production

Mechanisms of vocal production in songbirds are well understood and provide testable hypotheses for parrots. Songbird songs consist of sequences of discrete syllables separated by silent gaps, each containing rich subsyllabic acoustic structure^9^. Across songbird species, song is driven by a temporally precise neural sequence in the premotor nucleus HVC, where neurons fire sparsely at specific timesteps relative to syllable onset, forming a chain that tiles the whole syllable^15–17^. Critically, when subsyllabic acoustic elements are reused across distinct syllables, they are driven by distinct neural sequences in HVC^15–17^. HVC neurons thus encode the timestep within a syllable rather than a specific acoustic target. Even in juvenile birds, a given timestep within a differentiating syllable can specify distinct acoustic targets^15^. This code supports accurate imitation of a fixed number of syllables but limits subsyllabic flexibility and repertoire size because each additional syllable requires an independent neural sequence in HVC, regardless of acoustic similarity^15–17^. For example, an HVC-like code in Disco would be predicted to produce discrete neural sequences for ‘bird’ and ‘baby’ despite the shared ‘b’ sound, whereas an acoustically grounded vocal motor code would reactivate for the ‘b’ sounds independent of the acoustic context or time from syllable onset (Fig. 1a)^18,19^.

Parrots diverged from songbirds over 40 MYA and evolved a nucleated forebrain song system with more nuclei and connectivity (Fig. 3a-b), and its finescale projection patterns are unclear^20,21^. In the songbird song system, HVC is the only nucleus that projects non-topographically to the basal ganglia nucleus Area X (Fig. 3c), which is topographically aligned with downstream vocal motor cortical and brainstem vocal motor neurons^22^. HVC therefore broadcasts time step information across the full extent of Area X. In reinforcement learning terms, this architecture provides a ‘state space’ representation because each time step, treated as a state, can be flexibly mapped to the multiple vocal motor channels through synaptic plasticity^23^.

**Figure 3.**
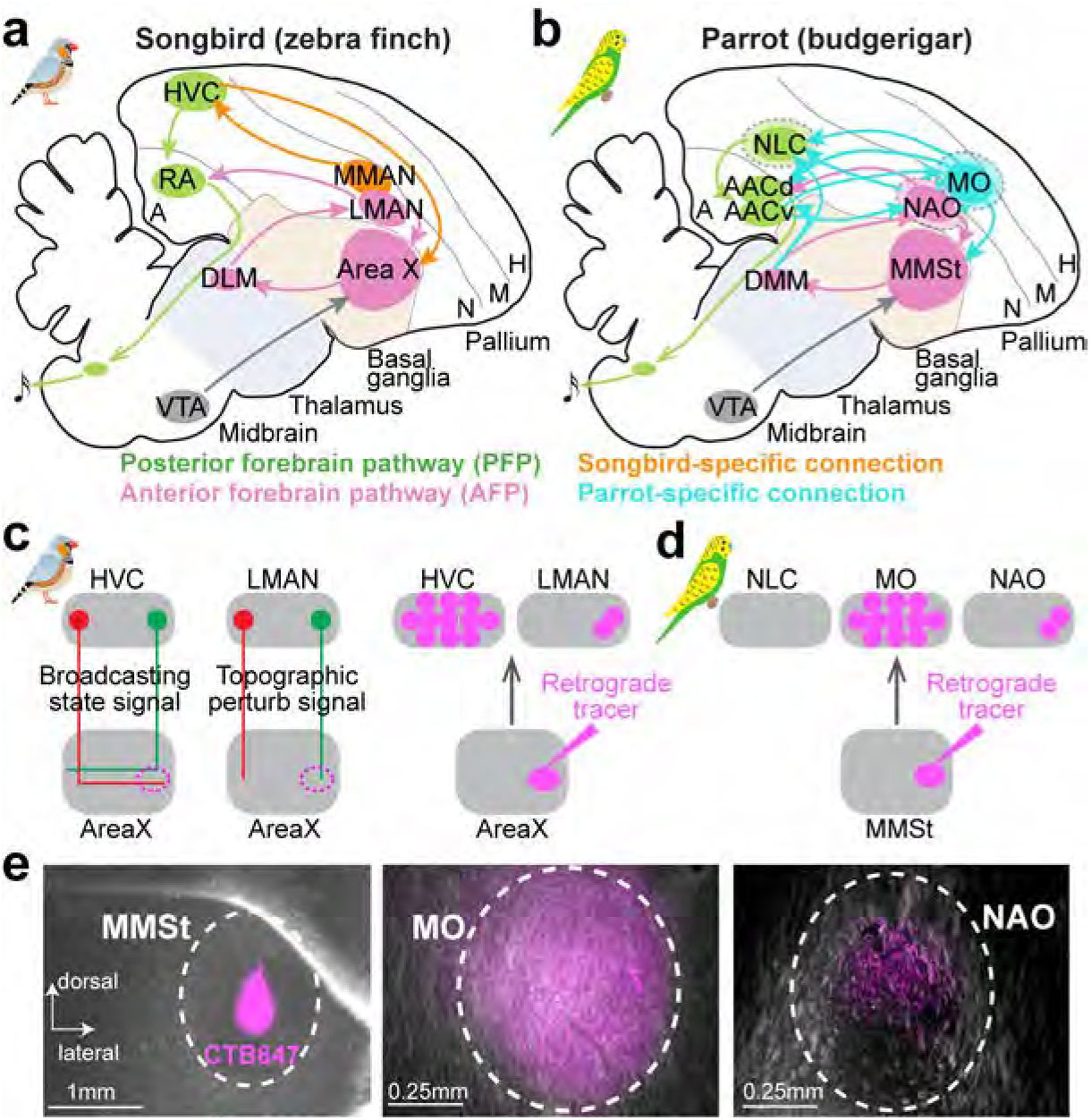
Inactivation and tracing experiments found that MO broadcasts song state signals. a-b, Diagrams of the vocal learning and production circuits in songbirds and parrots. For abbreviations, see legend for Extended Data Fig. 4c. c, Songbird reference results. Left, HVC broadcasts song state information broadly to the basal ganglia nucleus Area X, whereas LMAN provides topographically organized perturbation signals. Dots represent neuronal somata; lines represent axonal projections. Right, focal retrograde tracer injections into Area X backlabel neurons throughout HVC but only in spatially restricted regions of LMAN. d, Schematic summary of retrograde tracing experiments in parrots. e, Focal injection of retrograde tracers in parrot MMSt (left) backlabeled neurons throughout MO (middle), but only in restricted regions in NAO (right).

To find a parrot nucleus that could serve as a state space node for reinforcement learning, we looked for a region that projects non-topographically to the striatum. We injected small volumes of retrograde viral tracer into restricted regions of MMSt (the parrot analogue of songbird Area X). The only nucleus with widely distributed back-labeled neurons was the oval nucleus of the mesopallium (MO) (Fig. 3d-e, Extended Data Fig. 4, n=5 birds, 7 hemispheres), a nucleus with widespread projections to all other forebrain song system nuclei^21^ yet without a clear analogue in the songbird (but see ref^24^). Nucleus NLC, previously proposed to have an HVC-like function based on its posterior position^25^, did not project to MMSt. Previously, we reported that inactivating output nuclei of the anterior forebrain pathway (AFP), including MO and the adjacent NAO (oval nucleus of the anterior nidopallium), disrupted the learned acoustic structure in contact calls^14^. These inactivations also abolished warble song production (Extended Data Fig. 5), leading us to wonder what vocal motor codes may exist in MO during warble song.

### MO neurons exhibit sparse bursts locked to reused subsyllabic acoustic targets in calls

We implanted 16-channel moveable electrode arrays into MO and recorded single unit activity during singing (Extended Data Fig. 6a-b, Methods). We identified two main classes of neurons: sparse, temporally precise cells (n=241) and tonic, singing related cells (n=20) (Fig. 4a-b, Extended Data Fig. 6-7, Methods, n=4 birds, 5 hemispheres). The two classes differed significantly in sparseness index (mean ± SEM: 0.415 ± 0.010 vs. 0.036 ± 0.004, p < 10⁻⁴, Wilcoxon rank sum test) and baseline firing rate (3.56 ± 0.18 Hz vs. 42.63 ± 3.65 Hz, p < 10⁻⁴, Wilcoxon rank sum test).

**Figure 4.**
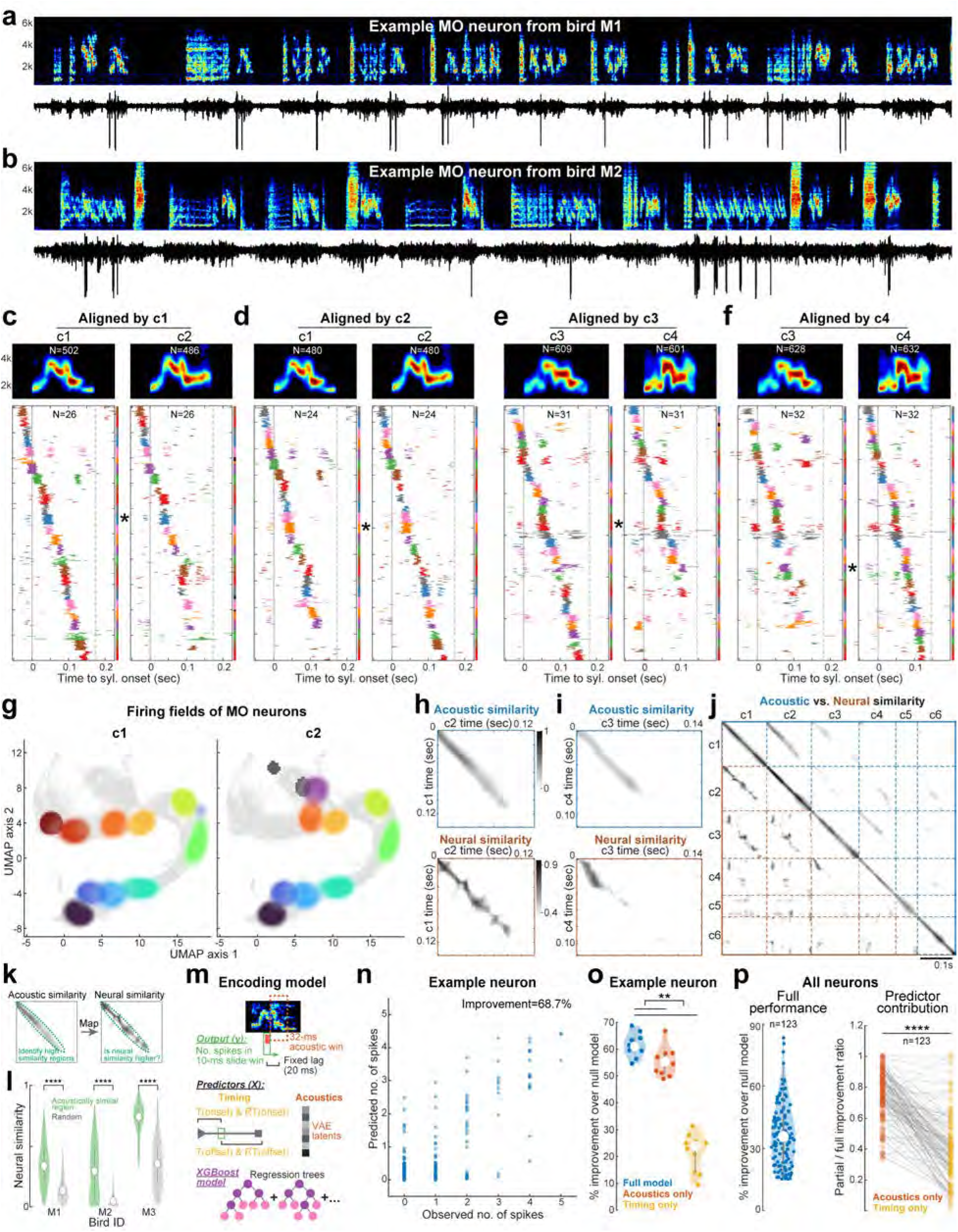
MO neurons form sparse and combinatorial sequences for producing warble calls. a-b, Spectrograms and voltage traces of two example MO sparse neurons. c-f, Bottom, population raster plots for sparse MO neurons that burst during the production of warble calls in bird M1, aligning to different call subtypes. Each row represents one rendition, with neurons colored differently using 8 colors that are reused periodically. In a given panel, the same color on the same row location indicates the same neuron. The colored bars on the right indicate the range of the renditions for a given neuron, with black bars indicating inadequate rendition numbers. Top, mean spectrograms of all call renditions that were used in the population raster plots below. Numbers indicate the total number of renditions. g, Firing fields of MO neurons (n=10) that bursted during the production of call c1 (left) and c2 (right), overlaid on the UMAP trajectories . The same color indicates the same neuron across panels. h, Relationship between acoustic and neural similarity for call subtypes c1 and c2. Top, acoustic similarity matrix. Bottom, neural similarity matrix computed from time-resolved SVM decoding of call identity (Methods). Regions of high acoustic similarity correspond to regions of high neural similarity. i, Same as in (h), but for comparison between call subtype c3 and c4. j, Summary across all pairwise call subtype comparisons in bird M1. Upper right triangle shows acoustic similarity (blue); lower left triangle shows neural similarity (brown). k, Schematics showing how summary statistics were calculated in (l). Regions with high acoustic similarity were first identified by thresholding, then mapped to the neural similarity matrix. We tested if regions with high acoustic similarity also have high neural similarity. l, Violin plots of neural similarity for regions with high acoustic similarity (green) and randomly selected regions (gray). The number of recorded MO neurons in bird M4 was inadequate for the pseudo-population analysis. Significance level was established with the Wilcoxon rank sum test. m, Schematics of the gradient-boosted decision tree model used to compare contributions of acoustic and timing predictors to MO spiking. n, Scatter plots of predicted vs observed number of spikes for an example neuron. Each dot indicates one sliding window. o, Model performance for the example neuron in n. Colored points indicate individual cross-validation folds; white points denote medians. Significance was assessed using Wilcoxon signed-rank tests. p, Population summary of model performance. Left, median performance for all neurons whose full models significantly outperformed null models (p < 0.05 and median improvement ≥ 15%). Right, relative contribution of acoustic and timing predictors, measured by the reduction in performance when each predictor class was removed from the full model. Each point represents one neuron; lines connect corresponding models. Statistical significance was assessed using Wilcoxon signed-rank tests.

We first examined activity of MO sparse neurons during the production of warble calls. Sparse neurons could discharge across multiple call types (mean ± SEM: 2.30 ± 0.12 call types across 4 birds). Aligning sparse neuron activity to distinct call types revealed precise neural sequences tiling each call, in apparent similarity to songbird HVC chains (Fig. 4c-f)^15,16^. In bird M1, neurons sorted by burst time during call c1 formed a clear temporal sequence spanning that call. Yet unlike HVC, the same population of neurons also precisely tiled the distinct syllable c2, but only for the first ∼100 ms of the call during which c2 and c1 shared a subsyllabic acoustic trajectory (Fig. 2e-f, Fig. 4c). Following acoustic divergence, neural sequences diverged as well (Fig. 4c). Re-sorting neurons by burst timing relative to c2 revealed a sequence of MO neurons that also fully tiled c2; a sequence that is shared with c1 only until the acoustic trajectories of c1 and c2 diverged (Fig. 4d). This trend generalized across call types and birds: shared acoustic trajectories corresponded to shared neural sequences, while distinct acoustics recruited distinct sequences (Fig. 4c–f, Extended Data Fig. 8). To visualize this relationship, we overlaid spikes from neurons active in c1 and c2 calls onto the UMAP projection of their acoustic trajectories. MO neurons discharged in consistent regions of acoustic space regardless of call identity, overlapping where acoustics overlapped and diverging where acoustics diverged (Fig. 4g).

We quantified neural-acoustic correspondence in three ways. First, support vector machine (SVM) decoders were trained to predict call type from neural activity in 10-ms sliding windows (Methods). If the SVM is unable to accurately distinguish two call types at a particular time point based on the neural activity, it indicates that those calls share a similar neural encoding at that time. Measuring decoding accuracy across all time points between two call types therefore yields a neural similarity matrix for each pair of call types (Methods). Comparing pairs that share acoustic segments, we found that neural similarity tracked acoustic similarity: neural decoding accuracy dropped during shared acoustic segments and rose when calls diverged (Fig. 4h-j, Extended Data Fig. 8). In all birds with sufficient neural data for SVM decoding analysis, regions with high acoustic similarity exhibit significantly higher neural similarity than overall background (n=19 pairs of call types over 3 birds, Fig. 4k-l). Second, we trained gradient boosted tree models^26^ to directly predict MO spikes from timing variables (absolute/relative latency to syllable onset/offset), VAE acoustic features, or both (Fig. 4m-n, Methods). We discovered that MO discharge was significantly better explained by acoustics than by timing (Fig. 4o-p). Finally, trial-by-trial variability in MO burst features (number of spikes in the burst) correlated significantly with conventional acoustic parameters such as pitch and amplitude with a lag of ∼25 ms, consistent with a premotor delay from MO to downstream motor regions (p < 0.001 for 57.5% of 299 neuron-call pairs, Extended Data Fig. 9). Notably, single MO neurons could be correlated with multiple acoustic metrics (Extended Data Fig. 9e-f).

Together, these results show that MO, like songbird HVC, contains sparse neural sequences that tile learned syllables. But whereas HVC neurons fire at fixed timesteps within a syllable and are not reused across syllable types, MO neurons form flexible sequences aligned to sub-syllabic acoustic targets that recur across call types.

### MO neurons encode acoustic targets regardless of timing and acoustic context in compound syllables

If MO neurons encode targets in an acoustic space, their discharge should be locked to specific region(s) of that space independent of the acoustic trajectory into or out of it, analogous to hippocampal place cells that discharge regardless of the path taken through their place fields^27^. Compound syllables exhibit highly variable acoustic structure across renditions (Fig. 5a, Extended Data Fig. 1), analogous to diverse paths in acoustic space, thus providing an ideal testbed for this hypothesis.

**Figure 5.**
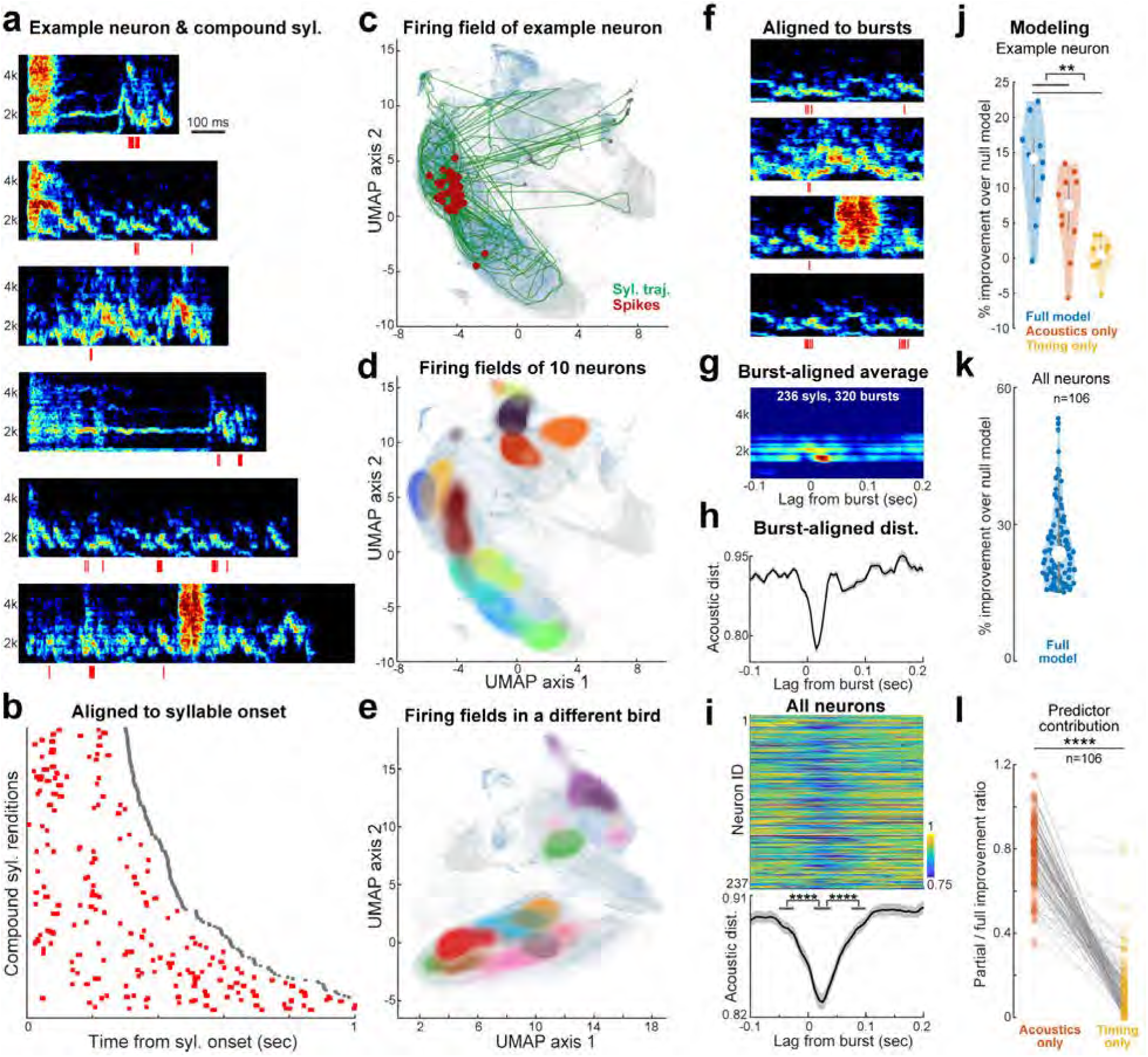
MO neurons encode acoustic targets in compound syllables. a, Example compound syllables with spikes from a single sparse MO neuron (red ticks), aligned to syllable onset. B, Raster plot of the same neuron as in a, aligned to compound syllable onset across renditions. c, Spikes from the example neuron (red dots) overlaid on the UMAP acoustic space. Green lines indicate compound syllable trajectories; grey triangles and squares mark trajectory onset and offset, respectively. Grey background shows the embedding of all syllables, while blue highlights compound syllables. d, Firing fields of 10 MO neurons. Each colored field indicates the density of all spikes for a given neuron. The dark red neuron is the example neuron in (a-c). e, Same as in (d), but for neurons from a different bird. Note that UMAP embeddings were constructed separately for each bird. f, Example spectrograms and spikes (red ticks) of the same neuron in (a-c), but aligned to the center of neural bursts. g, Burst-triggered averaged spectrogram for the sample neuron. h, Acoustic distance surrounding MO bursts for the example neuron. Black line and grey shading indicate the mean and SEM of pairwise acoustic distance across compound syllable renditions. i, Population summary across all sparse MO neurons. Top, heatmap showing changes in acoustic distance as a function of lag relative to neuronal bursts (each row is one neuron). Bottom, mean acoustic distance across neurons. Shaded regions indicate SEM. Statistical significance was assessed using Wilcoxon rank-sum tests. j, Gradient-boosted decision tree model performance for the example neuron shown above. Colored points represent individual cross-validation folds; white points denote medians. Significance was assessed using Wilcoxon signed-rank tests. k, Population performance for neurons whose full models significantly outperformed null models (p < 0.05 and median improvement ≥ 15%). Blue points indicate median performance for individual neurons; white point indicates the population median. l, Contribution of acoustic and timing predictors, measured by the reduction in model performance when each predictor class was removed from the full model. Each point represents one neuron; lines connect corresponding reduced models. Statistical significance was assessed using Wilcoxon signed-rank tests.

Within compound syllables, MO burst time relative to syllable onset was highly variable (Fig. 5a-b, mean coefficient of variation ± SEM: 0.866 ± 0.021 on latency of the first spike, n=229 neurons), resembling the variation in the timing of reused acoustic targets themselves (Fig. 2j) and inconsistent with an HVC-like timing code^15^. Compound syllables formed highly variable trajectories in the UMAP plot, as expected from their variable acoustic compositions (Fig. 5c). Yet spikes from the same MO neuron tended to form clusters at specific locations in the projected acoustic space (Fig. 5c), indicating consistent encoding of acoustic targets across different compound syllable renditions. Different MO neurons occupied distinct locations, suggesting encoding of different acoustic targets and tiling of the acoustic space with the MO neuronal population (Fig. 5d-e).

We quantified the relationship between MO bursts and acoustic targets in compound syllables in two ways. First, if MO discharge promotes production of specific acoustic targets in variable contexts, then bursts should precede convergent acoustic outcomes. We computed acoustic distance in windows spanning 100 ms before to 200 ms after burst onset in compound syllables. Acoustic distance sharply decreased ∼25 ms after bursting (Fig. 5f-i), consistent with a premotor delay (Extended Data Fig. 9), indicating that MO activity drives a stable acoustic target across variable acoustic contexts and latencies from syllable onset. Second, we formally tested the contribution of timing and acoustic predictors using gradient boosted tree models^26^ trained to predict MO spikes from either syllable timing variables, acoustic features, or both (Fig. 4m, Methods). Notably, acoustic predictors alone matched full-model performance, whereas timing contributed little for compound syllables (Fig. 5j-l). Together, these findings demonstrate that MO neurons encode an acoustically defined state space whose elements can be flexibly recruited across distinct vocal contexts.

### MO neuronal discharge across syllable categories

Most MO neurons were active across multiple syllable types (mean ± SEM: 2.13 ± 0.07 syllable types per neuron; Extended Data Fig. 10a), but exhibited varying degrees of generalization across syllable categories. One subset of neurons encoded shared acoustic targets across categories, showing overlapping firing fields for calls and compound syllables (Extended Data Fig. 10b–e). We quantified this generalization by training boosted models to predict MO spiking from VAE acoustic features on compound syllables, and then testing model performance on calls that were excluded from training, or vice versa. Approximately 15% of neurons (24/165 for compound-trained models tested on calls) and 10% of neurons (15/151 for call-trained models tested on compound syllables) performed significantly better than null models (p < 0.05, Wilcoxon signed-rank test; Extended Data Fig. 10f–g), indicating shared encoding of acoustic targets across syllable categories.

By contrast, many MO neurons exhibited heterogeneous specificity across syllable types. This heterogeneity was exemplified by neurons that showed precise, acoustically grounded responses during narrowband syllables such as calls, but also discharged during broadband syllables such as squawks or clicks (n = 75/241 neurons; Extended Data Fig. 10h–k). Thus, MO neurons could exhibit stable and precise firing fields within a syllable category, while encoding distinct firing fields across categories, conceptually analogous to the ability of hippocampal neurons to express distinct place selectivity across different environments^28^.

## Discussion

Our results show that in both imitated English and natural warble, budgerigars reuse discrete sounds in a manner analogous to phoneme reuse in human speech. We identify a sparse, acoustically grounded premotor code in the parrot frontal cortical nucleus MO, in which brief bursts specify subsyllabic acoustic elements independent of syllable timing, identity, or surrounding acoustic context. Individual MO neurons fire at specific regions of acoustic space across diverse vocal trajectories, acting as acoustic place cells whose activity is invariant to the path taken into or out of those regions.

Although both MO and the songbird premotor nucleus HVC generate sparse neural sequences that tile syllables, their coding strategies differ fundamentally. HVC neurons discharge at fixed timesteps within a syllable and are not reused when similar acoustic segments recur across syllables, forming a timing-based code^16,23,29^. Even during development, when two syllables transiently share an underlying HVC sequence, neuronal bursts remain locked to syllable time rather than ongoing acoustics^15^. This timing-based code supports accurate imitation of fixed acoustic templates^30^ but restricts subsyllabic flexibility: every new syllable requires a new neural sequence, even if it reuses existing acoustic building blocks^15^. Such mechanisms may support early acquisition of a fixed phoneme inventory in humans^9^, but they do not explain fluent recombination of units into the long vocal streams that characterize human speech^7^ (Fig. 1). In contrast, MO neurons represent acoustic targets directly: the same neuron can reactivate at the same acoustic region across times within a syllable and syllable categories. By reusing vocal motor codes within and across syllables, MO can support a large repertoire without requiring dedicated premotor chains for each gap-bounded syllable.

The acoustic place-like code in MO supports flexible sound production but raises the question of how vocal trajectories are selected and guided through acoustic space. MO activity predicted vocal output only within a brief premotor window of less than 50 ms (Fig. 5i), yet behavioral studies indicate that budgerigars can perceive and reproduce long, structured vocal sequences, including multisyllabic phrases^31^. MO receives extensive input from auditory areas^21,32,33^, which could transform desired vocal sequences or auditory targets into a series of MO activations that specify upcoming acoustic goals rather than immediate motor output. This organization raises the possibility that auditory inputs to MO may encode future vocal structure beyond the next acoustic element, which could be tested in future studies.

In addition to auditory inputs, MO projects to downstream motor regions, including the anterior arcopallium (AAC). Recent recordings from AAC identified neural population activity correlated with continuous acoustic features such as pitch and harmonic structure, and population activity could recur when acoustics recur within a single syllable^34^. Because AAC projects directly to brainstem vocal motor neurons^21^, these signals are well positioned to shape articulatory kinematics during sound production. Humans exhibit a broadly similar organization, in which reciprocal auditory-motor projections transform acoustic targets in auditory cortex into articulatory commands in precentral motor areas; recent intracortical recordings have identified neurons selective for individual phonemes or phoneme clusters during speech perception and productions^18,19,35^.

Together, these findings support a hierarchical transformation from sparse, reusable acoustic target representations to continuous articulatory control signals. In parrots, our results identify MO as a candidate locus for acoustic target representations upstream of AAC. In humans, although phoneme-selective neurons have been observed^18^, a sparse acoustic-target code with the degree of path invariance and context independence we observe in MO has not yet been identified. Our findings therefore motivate targeted searches in human recordings for sparse activations that may be difficult to detect with low-impedance electrode arrays and automated spike sorting.

In sum, we identify a premotor coding strategy in the parrot forebrain in which sparse, acoustically grounded bursts specify reusable vocal targets that can be flexibly recombined across contexts. This code supports large vocal repertoires without requiring distinct premotor sequences for each syllable, providing a mechanistic solution to the scaling problem of repertoire size. By revealing an explicit acoustic target representation in a circuit linking auditory cortical areas to primary vocal motor cortex, our findings motivate targeted searches for sparse phoneme-like codes in homologous regions of the human speech network.

## Acknowledgements

We thank members of the Goldberg lab for comments and Irene Pepperberg and Robert Dooling for inspiration. We thank John Pearson, Miles Martinez, Tim Sainburg, and Richard Hahnloser for discussion on acoustic analysis methods. This work was supported by funding to J.H.G. from National Institutes of Health (R34NS121898). Z.Z. was supported by the Klarman Fellowship from Cornell University and the Fellows-to-faculty Award from Simons Foundation.

## Author contributions

Z.Z. and J.H.G. designed the study. Z.Z. collected the data with help from C.J. and B.K.. Z.Z. and A.N.P. analyzed the data with help from C.J., B.K., and H.K.T.. J.H.G. and Z.Z. wrote the manuscript with help from all other authors.

## Materials and Methods

### Animal subjects and regulatory information

Animal care and experiments were carried out in accordance with NIH guidelines and were approved by the Cornell Institutional Animal Care and Use Committee. Subjects used in this study were 10 male and 3 female budgerigars (*Melopsittacus undulatus*) aged between 1 and 3 years. Birds were bred in our aviary or obtained from outside breeders (Magnolia Bird Farm, California; OK Birds, Oklahoma). We used 3 females and 2 males for anatomical tracing, 4 males for reversible inactivation, and 4 males for electrophysiological recording. Birds were first group-housed in flight cages before experiments, then housed in dyadic recording chambers for behavioral and neural recording. Food and water were provided ad libitum, with a 12-12 light-dark cycle. Colony noise was played during the light cycle with two computer speakers in the recording room.

### Behavioral recording for inactivation and electrophysiology experiments

Two birds were housed in acrylic dyadic chambers, with a transparent acrylic divider in between. A hole of diameter ∼0.8 inch in the center of the divider allowed birds to sing and interact with each other. Two one-way mirrors were placed on the side wall of each chamber, such that birds could also sing and interact with the mirror reflection. To record vocalizations, two unidirectional microphones (Pro 35, Audio-Technica) were placed on the central divider, pointing in the opposite direction into each chamber. Audio signals were first bandpass-filtered (30-15,000 Hz) and amplified with a preamplifier (XL48, Midas), then digitized to 50 kHz through a DAQ (USB-6210, NI) before sending to the computer. Videos were recorded at 90 fps for behavioral experiments and 60 fps for electrophysiology, using Blackfly cameras (BFS-U3-70S7C and BFS-U3-19S4M). We placed cameras behind the back wall of the dyadic chamber to record social interactions between two birds, or behind the one-way mirror to record interactions with the mirror reflection. Audios and videos were synchronized using a custom GUI in Python (https://github.com/GoldbergLab/PyVAQ). For electrophysiology experiments, a copy of audio signal and camera trigger signal was routed to the Intan recording controller (C3004) to synchronize with neural recordings sampled at 20 kHz. We calibrated the gain of each audio preamplifier channel such that differences in the sound amplitudes between two unidirectional microphones allowed reliable identification of the singing bird when only one bird was singing. For electrophysiology, the identity of the singing bird was further confirmed with head-mounted accelerometer data. When two birds were vocalizing simultaneously, data was excluded from further analysis.

### Tracing and histology

To perform surgery and histology in budgerigars, we adopted procedures detailed in Zhao et al. 2023. Stereotactic coordinates of MMSt, NLC, and AAC were first estimated based on the published budgerigar brain atlas (http://www.brauthlab.umd.edu/atlas.htm), then confirmed with tracer injections. We injected a small volume (9.2 nL) of tracers using Nano-ject II (Drummond Scientific), at the rate of 4.6 nL/pulse and 30 s interpulse intervals. For retrograde tracing, we used Cholera Toxin Subunit B (CTB) conjugated with Alexa fluorophores (Invitrogen, C34778) and self-complementary AAV viruses expressing EGFP or mRuby (ETH VVF, v65 or v286)^36^. One week post injection, animals were perfused and their brains were fixed in 4% paraformaldehyde solution for 1-2 days, then cut into 100 µm coronal slices with vibratome and mounted on slides in polyvinyl alcohol (Sigma-Aldrich, 10981-100ML). Dark-field and fluorescent images were acquired using an epi-fluorescence microscope (Leica DM4000 B) or a confocal microscope (Zeiss, LSM710).

### Reversible inactivation

We made custom microdialysis probes according to the method in Aronov et al. 2008. The inflow polyimide tubing is approximately 2.25 mm long for MO/NAO implant. The cellulose membrane (Replican, 132294) measures roughly 0.2 mm in width and 1 mm in length, with 0.7 mm exposed to deliver drugs. For MO/NAO, the probes were implanted bilaterally at 8.80 AP, 2.90 ML and 1.45 DV (unit: mm), with a 30° AP angle and 20° ML angle. To secure the implants, we inserted 4 insect pins into the skull and covered the base of probes with dental cement (C&B Metabond, Parkell, S380). The probes were flushed daily with PBS to prevent clogging. After data collection, we infused fluorescein (10 mM) into the probe to confirm implant location and drug diffusion.

Compared to direct injection, a higher concentration of drug is needed with the microdialysis probe, since the solute needs to diffuse passively through the cellulose membrane ^37^. Following previous studies in zebra finches ^37^ and budgerigars ^14^, we diluted tetrodotoxin citrate (Cayman, CAYM-14964-1) to 50 µM and muscimol (VWR, Q-1345.0050BA) to 1.5 mg/mL in PBS. One week after implant surgery, we started infusing PBS into probes to acclimate birds to the procedure. Infusion was conducted ∼1 hour before the cage light turned on. For each probe, we infused ∼0.5 mL solution at a rate of ∼0.1 mL/min. The bird was then placed back to the behavioral chamber to rest before the light turned on. After ∼6 hours of behavioral recording, the drug was washed out with PBS. We used the same procedure for drugs and PBS, and conditions were alternated daily, so a direct comparison is meaningful. We had 2-3 infusion replicates for each bird under each condition.

### Electrophysiological recording

#### Surgery

For chronic neural recordings, birds were anesthetized with isoflurane inhalation and mounted on a stereotaxic instrument for probe implantation. All probes were 16-channel moveable electrode bundles (Innovative Neurophysiology). The anterior skull was first leveled, then the head was adjusted to a 30° AP angle. Probes were implanted at AP 8.80, ML 2.50, and DV 0.50 (unit: mm), relative to the sagittal sinus. To secure the implant, we inserted 4 insect pins into the skull and covered the base of probes with dental acrylic (Pentron Clinical, N11H). Three birds were implanted unilaterally at the right hemisphere, while one bird was implanted bilaterally. Animals were allowed to fully recover from surgery before recording started.

#### Recording

We extended or retracted (50 or 100 µm) electrodes on a semi-daily basis by manual turning of a screw in the morning before the enclosure lights turned on. To confirm the recording location, we took two approaches: 1) we dyed the electrode bundle with Dil (ThermoFisher, D282) before the surgery and visualized the dye location using a fluorescence scope, and 2) we burned selected electrode channels that had sparsely bursting neurons by passing a small current (30 µA) for 30 s after the recording experiment was completed, then examined the burned location under a microscope. Neural signals were acquired with the Intan recording controller (C3004) and Intan headstages (RHD2132) that directly interfaced with the moveable bundles. Sampling rate was set to 20 kHz and recording was manually controlled with the Intan recording software, where a 60 Hz notch filter was applied and spiking activity of single units could be visualized in real time. The Intan headstage is equipped with a 3-axis accelerometer that samples at 5 kHz.

#### Spike sorting

A custom MATLAB GUI was used for visualizing vocalization and neural data, and for spike sorting. Neural recordings were bandpass filtered between 400 Hz and 9000 Hz, and single unit spiking activity was manually sorted as previously described ^38^. We sorted neurons with signal/noise ratio larger than 10 between the dorsal-ventral range 0.5-1.8 mm, where MO is located and the majority of sparsely bursty neurons were identified.

### Vocalization detection and warble identification

We automatically detect vocal activity by first calculating spectral flatness of the recorded audio stream, then identifying candidate vocalizations by using the findpeaks function in Matlab with a threshold of −0.75. To automatically identify warble songs, we first adopted a set of relaxed criteria to search for candidate vocal sequences that 1) had at least 5 syllables; 2) gaps between syllables smaller than 10 s; 3) total duration longer than 5 s; 4) peak syllable density (number of syllables in a 5 s sliding window) larger than 3. We then manually examined candidate vocal sequences and excluded sequences composed of 1) pure cage or movement noises; 2) pure contact calls or squawks; 3) two birds vocalizing simultaneously. The remaining sequences were marked for spike sorting.

### Analysis of Disco vocalizations

Disco data source:

I’m Disco and I know it: https://www.youtube.com/watch?v=EFJeY9fL5tk

A splattered modern major doo: https://www.youtube.com/watch?v=Wmpep__aTO4

Disco the parakeet chats and mumbles: https://www.youtube.com/watch?v=2X4CTOC8LF0

I like history and call me Shirley: https://www.youtube.com/watch?v=KXuaOvxI55g

I’m a parakitty: https://www.youtube.com/watch?v=xhtme--h5Zs

nobody expects a bird to quote monty python: https://www.youtube.com/watch?v=hNavxvvN9QI

One phrase leads to another: https://www.youtube.com/watch?v=WYcWA_RNl1s

That’s a Discolicious burger: https://www.youtube.com/watch?v=62zUKOUETVY

The rules by Disco the parakeet: https://www.youtube.com/watch?v=OavCadkEycU

OpenAI’s Whisper model and a human listener independently annotated 26 minutes of raw audio from Disco. Words agreed upon by both annotators were converted to phoneme strings using a text based grapheme to phoneme (G2P) tool. Vocalizations were segmented from silence using amplitude thresholding followed by manual curation. Using the same procedure described in Acoustic-space construction below, we trained a shotgun variational autoencoder (VAE) on 32 ms spectrogram windows extracted from segmented vocalizations with a 1 ms hop. To identify short within syllable acoustic segments that were reused across the repertoire, we used a single utterance of the word “Spanish” as a reference. “Spanish” was chosen as a representative multisyllabic word containing diverse phoneme-like elements; this analysis was not intended to exhaustively inventory all elements. We computed the cosine distance matrix between the VAE latent trajectory of this reference utterance and the latent trajectories of all segmented vocalizations in the 26 minute dataset. High similarity matches were defined as continuous strands in this matrix with cosine distance ≤ 0.15 sustained for ≥ 20 ms. To identify frequently reused segments within the reference utterance, we computed, for each reference time point, the probability of participating in a high similarity strand across the dataset and used peak detection to define candidate “acoustic primitives.” Here and in the primitive identification analyses below, cosine distance thresholds used to define high similarity strands should be interpreted as conservative, user defined operating criteria rather than estimates of a natural clustering boundary in latent space. Qualitative conclusions were unchanged across a wide range of thresholds (e.g., cosine ≤0.1–0.35; duration ≥15–50 ms).

### Warble segmentation and annotation

Previous studies^12,39^ used two paradigms to classify syllables in the complex budgerigar warble song: 1) elemental or compound syllables according to composition and complexity; 2) narrowband, broadband, or harmonic syllables according to frequency modulation pattern.

Built upon these efforts, we classified warble syllables into five broad categories to facilitate neural analysis: 1) calls, narrowband elemental syllables, with frequency band centered ∼3 kHz and duration ∼200 ms; 2) clicks, broadband elemental syllables, with noise-like frequency pattern, duration ∼40 ms, and soft sound amplitude; 3) squawks, broadband elemental syllables, with noise-like frequency pattern, duration ∼120 ms, and loud sound amplitude; 4) chortles, long compound syllables, with at least two repetitive narrowband acoustic units that are centered around 2 kHz; 5) others, all other elemental and compound syllables that are not categories 1-4. Together, categories 1-4 account for 60-70% of all syllables in the warble of recorded birds. Example spectrograms of each syllable category are shown in Extended Data Fig. 1a.

We fine-tuned WhisperSeg-base, a generic transformer model trained on animal vocalizations ^13^, to automatically segment warble syllables from silence and classify syllables into these 5 broad categories. We adopted an iterative and bootstrapping approach for the fine-tuning process. First, we segmented a small dataset (4 birds in a separate behavioral study, total 25 min of warble and 5916 syllables) by amplitude thresholding with manual adjustments and manually annotated syllables into the 5 broad categories using a custom Matlab GUI. Second, we fine-tuned the WhisperSeg-base model on this dataset for 5000 iterations using the train.py script in the WhisperSeg package with default parameters. Third, we used the trained model to predict syllable boundaries and labels on an expanded dataset (5 birds in current study). Fourth, we manually corrected the predicted syllable boundaries and labels to generate an expanded training dataset (62.5 min of warble, 9379 syllables). Fifth, we fine-tuned WhisperSeg-base on this expanded dataset from scratch for 5000 iterations using default parameters, to generate a final model. To evaluate the accuracy of this model, we manually segmented and labeled an independent test dataset (5 birds in current study, 10 min of warble) that were never used in model fine-tuning, and calculated F1 scores by comparing predicted labels with hand labels. Overall, the model achieves satisfactory accuracy with an averaged F1 score of 0.878 (Extended Data Fig. 1b).

### Warble call clustering and subtype identification

To facilitate alignment of neural responses, we further categorized call syllables into subtypes using unsupervised clustering. We adopt a deep learning-based pipeline that was shown to achieve state-of-the-art clustering accuracy over multiple animal vocalization datasets ^3,10,40,41^. First, annotated call syllables from WhisperSeg analysis were bandpass filtered (250-7500 Hz) to remove low- and high-frequency noise, then converted to spectrogram images with the stft function in the *scipy* python package (12.8 ms window size and 4 ms sliding length). Second, we embedded each call spectrogram at the center of a 128*128 matrix, without stretching on the time or frequency axis. The longest time axis was chosen to be the 0.995 quantile of the duration distribution, rather than the max duration, to prevent the reduction in temporal resolution caused by WhisperSeg misannotating long compound syllables as calls. Syllables longer than the duration threshold were clipped off at both edges. Third, we trained a variational auto-encoder (VAE) to reduce the 128*128 spectrogram images to 32-dimensional latent features that best capture variability in the spectrogram data^10,14^. We used the same VAE architecture as in Goffinet et al 2021, which consisted of 7 convolutional blocks and 4 dense blocks for the encoder, a 32-dimensional layer for the bottleneck, and 4 dense blocks and 7 transpose convolution blocks for the decoder. The network was trained on the entire call dataset of a bird for 51 epochs with batch size 64 and learning rate 0.001. Fourth, we performed UMAP (Uniform Manifold Approximation and Projection) using the *umap* python package on the 32-dimensional latent features to obtain 2D embeddings for data visualization and clustering (parameters: n_components 2, n_neighbors 25, min_dist 0, distance metric cosine). Fifth, we conducted clustering using HDBSCAN (Hierarchical Density-Based Spatial Clustering of Applications with Noise) using the *hdbscan* python package on the UMAP embeddings (parameters: min_samples 20, cluster_selection_method ‘leaf’, min_cluster_size ∼5% of the total sample size). Data from different birds were analyzed separately to maximize the separation between different call subtypes.

To confirm that clustering is not an artifact of the nonlinearity caused by UMAP or HDBSCAN, we performed control analysis as outlined in ref^10^. Briefly, we randomly sampled 32-dimensional latent features from a multivariate Gaussian distribution with mean and covariance matrix matched to the trained VAE latent distribution and sample size matched to the real dataset. We then performed the UMAP/HDBSCAN analysis on the randomly sampled dataset using the same parameters. We compared clustering quality between the real and random datasets using the Silhouette score in the UMAP space and Calinski-Harabasz index in the original VAE space and UMAP space^3,10^. Unlike the Calinski-Harabasz index that remains robust in high-dimensional space, Silhouette score is sensitive to high-dimensional sparsity (the curse of dimensionality)^42^, thus not used for comparison in the original VAE space, although the calculated values are still higher for the real dataset than the random dataset. Random sampling was performed for 200 rounds to obtain a null distribution of the clustering quality metrics.

The quality of clustering varied across call subtypes. To identify low-quality clusters, we reasoned that calls have narrow frequency bands and well-defined call subtypes should have stereotyped frequency contour. We first time-warped all renditions of a given call cluster and calculated spectrograms that were aligned on the temporal axis. We then calculated an entropy-based sparseness index (for a definition see *Syllable-level analysis on neural activity*) on the frequency axis at each time point of the call. The median sparseness of all time points was then used as a quality metric for the call cluster. Finally, we applied a threshold (0.12) and excluded clusters with lower sparseness index from further acoustic and neural analysis.

### Acoustic-space construction

#### VAE training

We adopt a “shotgun VAE” approach to construct a low-dimensional acoustic space for visualizing and analyzing the warble song and neuronal activity^3,10^. First, all annotated syllables from WhisperSeg analysis were bandpass filtered (250-7500 Hz) to remove low- and high-frequency noise, then converted to spectrogram images with the stft function in the scipy python package (12.8 ms window size and 1 ms sliding length). To analyze pre-onset and post-offset activity, we also included 32 ms (duration of one sliding window in the second step below) of the silent gaps before syllable onset and after syllable offset. If the gap between the focal syllable and upstream/downstream syllables was shorter than 32 ms, additional zeros were padded. Second, we slid a fixed-duration window (32 ms) over the syllable spectrograms to create a training dataset for VAE. We randomly selected spectrogram windows from each syllable with sample size proportional to the syllable duration, i.e. more windows from longer syllables. Each selected spectrogram window was then interpolated to 128x128 pixels. The total number of spectrogram windows in the training dataset is ∼200k for each bird. Third, we trained a VAE network with a 32-dimensional bottleneck between the encoder and decoder for 51 epochs with batch size 64 and learning rate 0.001 ^10^. Fourth, after the VAE was trained on the random dataset, to obtain trajectory of a given syllable, we slid the window systematically from 32 ms before syllable onset to 32 ms after syllable offset with hop length 1 ms, then applied the trained VAE to obtain latent features of all spectrogram windows of this syllable. The resulting 32-dimensional latent features were used for downstream UMAP and decoding analyses.

#### UMAP

The 32-dimensional VAE latent features of sliding spectrogram windows were used as inputs for UMAP embedding. The following parameters were used: n_components 2, n_neighbors 25, min_dist 0, distance metric euclidean.

#### Acoustic distance in the VAE latent space

To compare acoustic similarity between two spectrogram sliding windows (not distributions) in the VAE latent space, we calculated the cosine distance between the two 32-dimensional vectors. Note that UMAP was used only for visualization, acoustic distance was calculated over the VAE latent vectors. For a group of sliding windows, we randomly selected 1000 pairs of sliding windows and averaged their cosine distances as a metric for the pairwise acoustic distance. The random selection process was repeated 20-50 times to obtain a distribution of the pairwise acoustic distance for the group.

### Identify primitives reused by calls and compound syllables

#### Identify compound syllables

We identified candidate compound syllables from the WhisperSeg categories ‘b’ and ‘x’ by setting a duration threshold (≥0.3 s). The duration threshold automatically excluded calls, clicks, and squawks that were mislabeled as ‘x’.

#### Discover primitives reused by calls and compound syllables

We selected a call subtype as the reference and calculated the acoustic distance matrix between the VAE latents of every candidate compound syllable and the mean VAE latents of the reference call subtype. We then identified strands with high acoustic similarity in the cosine distance matrix by setting a threshold on the distance (<0.2) and duration (≥20 ms). To identify tentative primitives that were used frequently between calls and compound syllables, we calculated the probability of having a high-similarity strands at each moment of the reference call and used peak detection to define the boundaries of the tentative acoustic primitives. The analysis was then looped over all call subtypes in all birds as the reference template and all compound syllables in the same birds.

### Syllable-level analysis on neural activity

#### Aligning spikes to syllable types

We aligned spikes from different MO neurons to warble syllable types, e.g. call subtypes and squawks, using methods from previous songbird studies^15,16,29^. For a given syllable type, we calculated a warping ratio for each rendition to the mean duration of all renditions, then linearly time-warped the audio data and spike times by the warping ratio. We then calculated neuronal firing metrics using the time-warped spikes. Given the premotor nature of MO, firing metrics were calculated over the range of 50 ms before the syllable onset to 0 ms after the syllable offset.

#### Inter-rendition correlation coefficient (IRCC)

To quantify rendition-to-rendition consistency in neural activity on short time scales, we computed the pairwise correlation coefficient of the instantaneous firing rate (IFR) between time-warped renditions of the same syllable type ^38^. For a given rendition, we first calculated the smoothed IFR by convolving the spike train with a Gaussian kernel of 10 ms standard deviation, then calculated Pearson correlation coefficients between all rendition pairs. Correlation coefficients from all rendition pairs and syllable types were then averaged to a single IRCC value for each neuron. To be included in the averaging, a neuron-syllable type pair needed to have at least 15 renditions and 25% of all renditions that have at least one spike within the range of 50 ms before the syllable onset to 0 ms after the syllable offset.

#### Sparseness index

To quantify the selectivity of neuronal activity during the production of a specific syllable type, we calculated an entropy-based measure of sparseness ^38,43^. For each neuron-syllable type pair, we first calculated the syllable-aligned PSTH (10 ms bin size, resulting in a total of *N* bins), which was normalized to firing probability vector *p* such that the sum of all values in this vector was equal to 1. We then computed an entropy-based sparseness index as 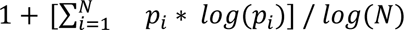 The sparseness index is equal to 1 (maximal sparseness) when activity is restricted to a single time bin of the syllable and goes to zero when spikes are evenly distributed throughout the syllable. Sparseness indexes from all syllable types were then averaged to a single value for each neuron. To be included in the averaging, a neuron-syllable type pair needed to have at least 15 renditions and 25% of all renditions that have at least one spike within the range of 50 ms before the syllable onset to 0 ms after the syllable offset.

#### Population raster plot

We selected sparse neurons to include in the population raster plots using four criteria on their firing metrics: 1) peak PSTH larger than 10 spikes/s; 2) at least 7 renditions and 20% of all renditions have spikes within 15 ms of the PSTH peak; 3) IRCC larger than 0.2; 4) sparseness index larger than 0.15. Different neurons are positioned according to locations of their peak PSTH values. As in Okubo et al. 2015, due to variability in the number of spikes for a given neuron-syllable pair, we constructed a population raster plot by choosing 20 renditions for which the neuron was most active around the PSTH peak. Note that this selection procedure is only for visualization purposes. Neuronal firing metrics and decoding analysis were calculated over all renditions. We confirmed that random selection of renditions resulted in noisier yet qualitatively similar population raster plots.

#### Correlating spike numbers with conventional acoustic metrics

To test if variation in neural activity is associated with variation in acoustics across diverse lags, we focused on calls and performed correlation tests between the number of spikes in MO bursts and conventional acoustic metrics. First, for a given neuron-call pair, we time-warped spikes to the mean duration of the call subtype. Second, we identified the time of an MO burst by finding the peak in the PSTH curve. Third, we counted the number of spikes within ±15 ms of the PSTH peak. Fourth, we calculated eight conventional acoustic metrics (Wiener entropy, pitch, goodness of pitch, spectral center of gravity, spectral width, amplitude, amplitude modulation, frequency modulation) using 9 ms windows with 1 ms slide ^15^. Fifth, we performed Spearman correlation tests between the number of spikes in the burst and the acoustic metrics with different lags.

### Firing fields of MO neurons in the acoustic space

We first constructed the acoustic space of each bird by running the shotgun VAE and UMAP pipeline with all syllable types combined (sliding window size 32 ms, hop 1 ms). For a given MO neuron, we overlaid spikes in the UMAP space by assigning each spike to the nearest sliding window with a premotor lag of 5 ms. The latency between the spike and the center of spectrogram sliding window is then 21 ms, which is roughly the latency with the highest correlation in the spike number-acoustic metrics analysis (Extended Data Fig. 9) and the lowest acoustic distance in the burst-aligned analysis (Fig. 5f-i). To calculate spike density, we divided the UMAP 2D space into a 100x100 grid and estimated density using a Gaussian kernel. The firing fields of multiple MO neurons were then visualized using different colormaps.

### Acoustic and neural similarity matrices

#### MMD and acoustic similarity index for calls

Maximum mean discrepancy (MMD) was used to quantify the distance between VAE latent distributions of two call subtypes ^10^. Essentially, MMD measures the distance between the mean embeddings of two distributions in a reproducing kernel Hilbert space. Following ref^10^, we used a Gaussian kernel and chose its width (sigma) to be ∼25% of the median distance between points in the aggregate sample in order to focus on finer differences in distributions. We varied the choice of sigma values and obtained qualitatively similar results. To obtain the MMD matrix between call subtype pair A-B, we first time-warped all renditions of A and B separately to their mean duration and interpolated VAE latents correspondingly, then calculated MMD between every timepoint x in subtype A and timepoint y in subtype B. For visual clarity, we decomposed the MMD matrix into a rank-1 matrix and a residual matrix (dMMD). The calculated dMMD values were further linearly transformed to acoustic similarity indexes by min-max normalization: (Vmax-dMMD) / (Vmax-Vmin) to the [0, 1] range, where low dMMD was transformed to high acoustic similarity index.

#### Neural similarity matrix between pairs of calls

To quantify the similarity structure of MO activity across call subtypes, we trained support vector machine (SVM) classifiers to decode call subtype identity from MO spiking activity. For each call subtype pair A–B, we randomly sampled renditions (n = 1000) from neurons that exhibited sparse bursts during the production of either A or B, forming a pseudo-population dataset. For each rendition, we computed instantaneous firing rates by counting spikes in a 10 ms sliding window with a 2 ms hop. At each time point, neural activity was thus represented as an N-dimensional vector, where N is the number of MO neurons included. For each pair of time points (tA, tB), we trained linear-kernel SVM models to discriminate between call subtype A at time tA and call subtype B at time tB. Models were trained on half of the dataset and evaluated on the held-out half. Decoding accuracy on the test set was used as a measure of neural dissimilarity between time tA of call A and time tB of call B. Repeating this procedure across all time point pairs generated a dissimilarity matrix D. This matrix was linearly transformed into a neural similarity index by min–max normalization: (Vmax-D) / (Vmax-Vmin) to the [0, 1] range. Here, Vmax=1 (perfect discrimination), while Vmin=0.5 (chance-level).

#### Statistical test on the relationship between acoustic similarity and neural similarity

For each pair of call subtypes, we first identified regions of high similarity in the acoustic similarity matrix by applying a threshold to dMMD values. This threshold was chosen to isolate strongly similar acoustic regions, rather than to define discrete acoustic categories. To focus on extended acoustic features, we further imposed a size threshold (≥ 50 matrix entries) to exclude small, isolated regions. These acoustically similar regions were then mapped onto the neural similarity matrix. Finally, we compared the distribution of neural similarity indices within these regions to those obtained from randomly sampled matrix entries using a Wilcoxon rank-sum test.

### Gradient-boosted decision tree models

#### Model inputs and outputs

We trained gradient-boosted decision tree models (XGBoost)^26^ to predict the number of MO spikes in 10 ms sliding windows (5 ms overlap). Two classes of predictors were included: timing and acoustic (Fig. 3m). Timing predictors (n = 4) comprised absolute latency from the spike window to syllable onset and offset, and relative latency to onset and offset (absolute latency divided by syllable duration). Acoustic predictors consisted of the 32-dimensional VAE latent vector corresponding to a 32 ms spectrogram window offset by a fixed premotor delay of 20 ms relative to the spike window (Fig. 3m). Gradient boosting was chosen because it captures nonlinear interactions between timing and acoustic predictors while remaining robust to correlated input features.

#### Model training

For each MO neuron, data were partitioned into 10 folds using syllable identity as the grouping variable, such that all data from a given syllable were assigned exclusively to either the training or testing set, preventing data leakage. At each iteration, one fold was held out for testing and the remaining nine folds were used for training. Models were trained using the XGBoost implementation in Python with the following parameters: objective = ‘reg:squarederror’, n_estimators = 200, max_depth = 10, learning_rate = 0.1, subsample = 0.9, colsample_bytree = 0.8, min_child_weight = 10, reg_alpha = 0.1, reg_lambda = 5, gamma = 1, random_state = 42, tree_method = “hist”.

#### Model evaluation

For each fold, model performance was evaluated by computing mean squared error on the held-out test set and compared to a neuron-specific null model that predicted the mean spike count across all inputs. Model performance was quantified as the percentage improvement over the null model, calculated as (error_null - error_model) / (error_null) * 100%.

## Code availability

All code and scripts used for data analysis were publicly available from the GitHub repository: https://github.com/zhileiz1992/MatlabCodesForZhao2025

---

**Extended Data Figure 1.**
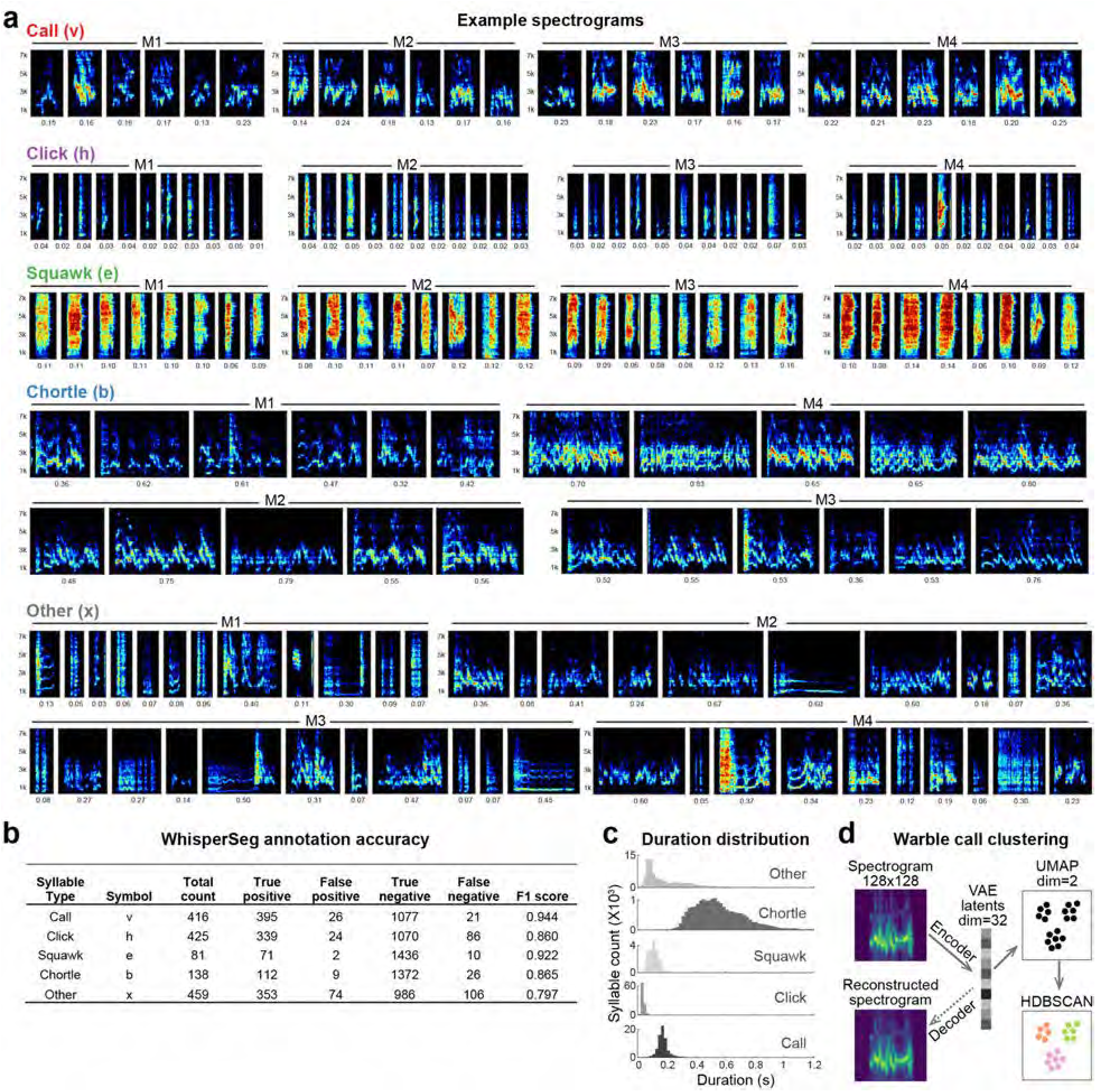
Example results and accuracy of WhisperSeg segmentation and annotation of budgerigar warble songs. a, Example spectrograms of the syllable categories labeled by WhisperSeg. Renditions were randomly selected from all WhisperSeg labeled syllables. The number below each spectrogram indicates syllable duration. Approximately equal number of renditions were shown for each bird. b, Accuracy of WhisperSeg syllable annotation evaluated on an independent, manually annotated test dataset. c, Distribution of syllable durations for each category. Shades of gray as in Figure 2b. d, Schematic of call clustering pipeline. VAE (variational autoencoder); UMAP (uniform manifold approximation and projection); HDBSCAN (hierarchical density-based spatial clustering of applications with noise).

**Extended Data Figure 2.**
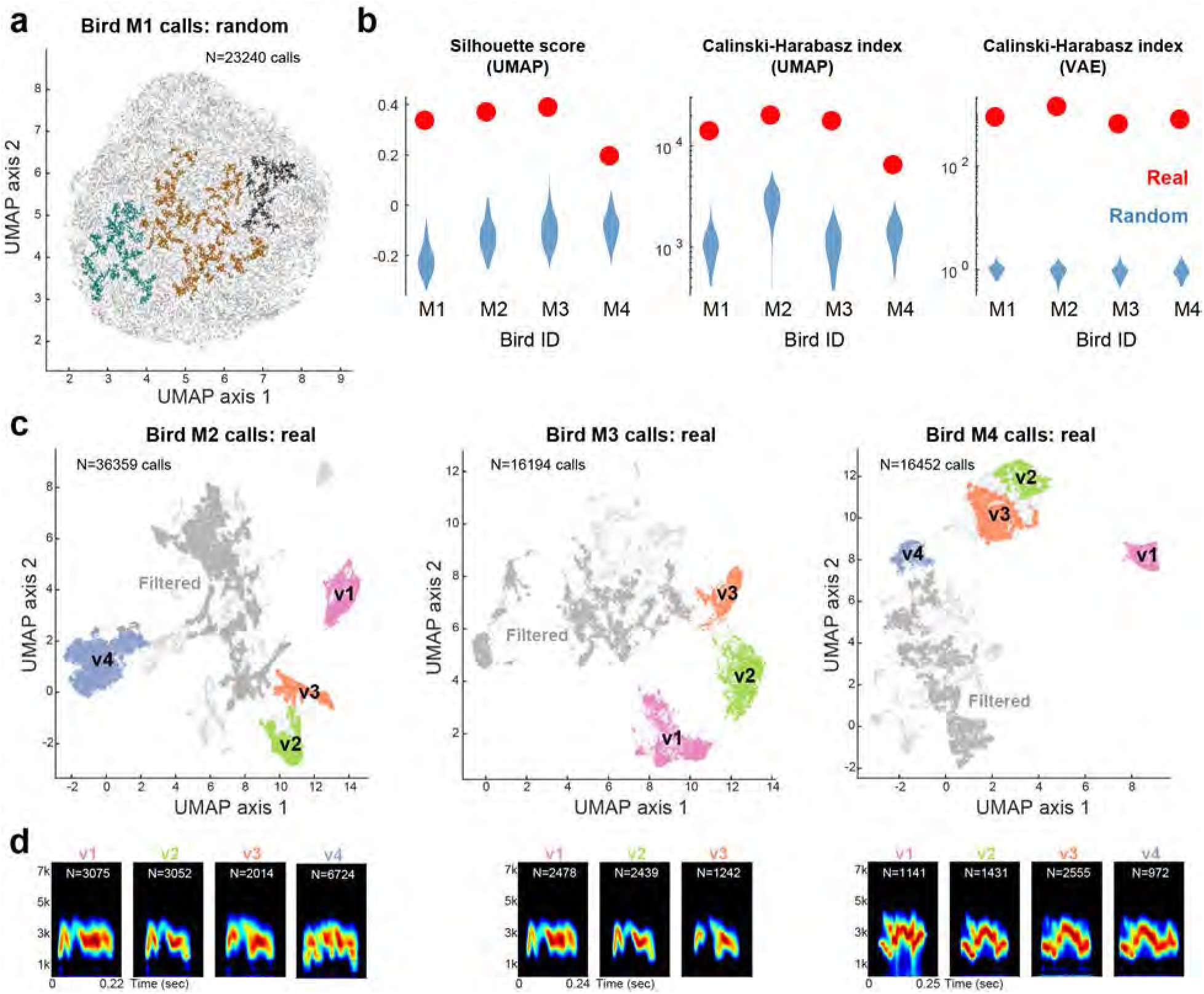
Clustering of warble calls from other birds. a, Control clustering analysis for bird M1. UMAP projection and HDBSCAN labels of a randomly generated call dataset drawn from a multivariate Gaussian matched to the VAE latent distribution of the real data (Methods). Each point represents one call; colors denote HDBSCAN cluster labels. Light grey indicates calls unclustered by HDBSCAN. b, Distribution of three clustering quality metrics (Methods). Clustering quality is significantly higher for the real datasets (red points) than for randomly generated datasets (blue; 200 randomizations). Because the Silhouette score relies on point-to-point distances, it is unreliable in high-dimensional spaces where distances concentrate; therefore, it is not used to evaluate clustering in the original VAE latent space, whereas the Calinski–Harabasz index remains robust. c, UMAP projection and HDBSCAN labels of warble calls for bird M2-M4. Each point indicates one call, colored by the HDBSCAN cluster label. Light grey indicates calls that were unclustered by HDBSCAN, while dark gray indicates call clusters that failed to pass our quality control procedure (Methods). d, Averaged spectrograms of call subtypes identified by HDBSCAN in (c) for bird M2-M4. Numbers inside the spectrogram indicate the total number of renditions identified and used for averaging.

**Extended Data Figure 3.**
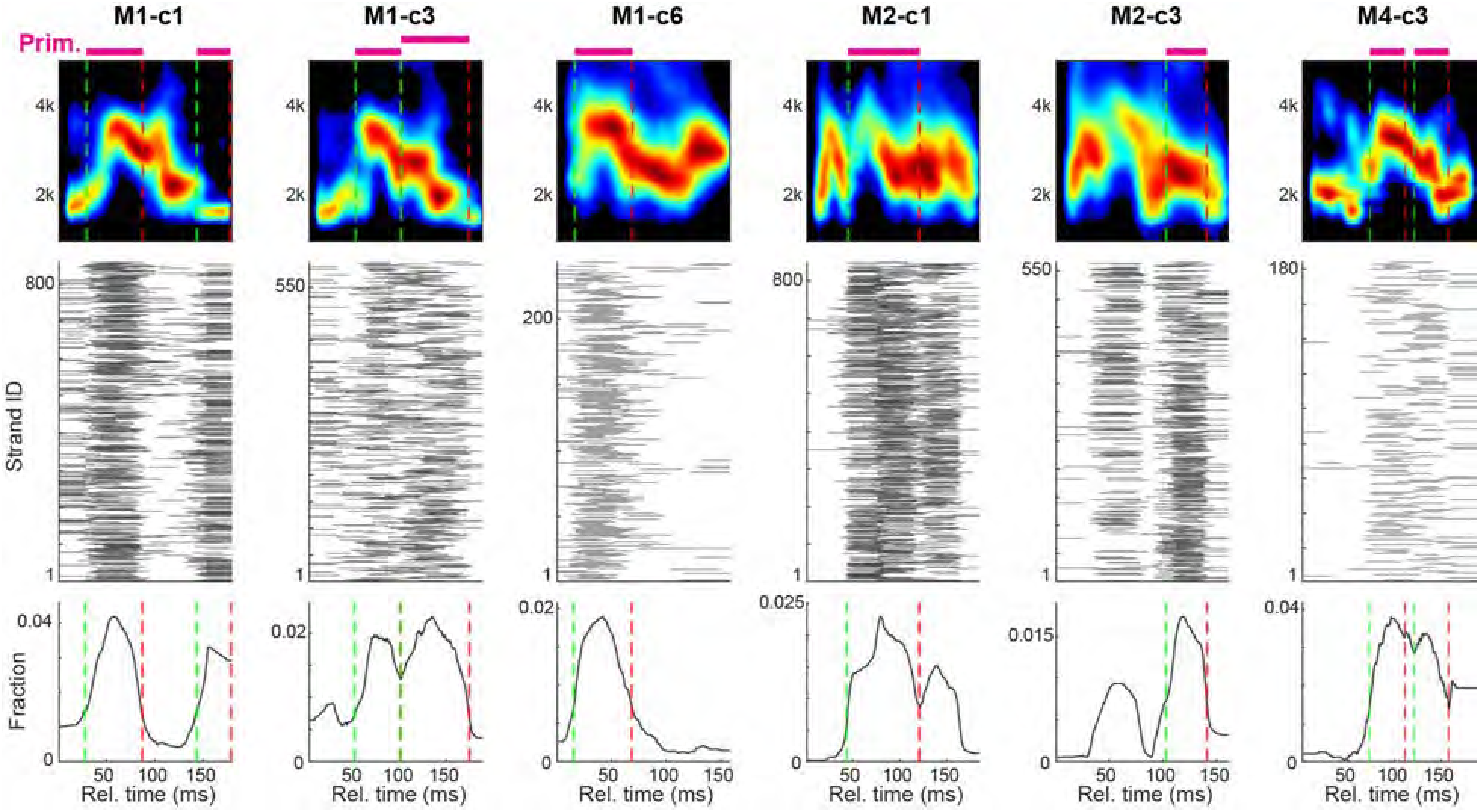
Additional examples of shared primitives between calls and compound syllables. Top, mean spectrograms of reference call subtypes. Middle, high-similarity strands from call–compound comparisons, plotted in reference time coordinates (each row represents one strand, as in Fig. 2i). Bottom, strand density across reference time. Peaks in strand density (dashed lines) define reused acoustic primitives, indicated by pink bars above the call spectrogram.

**Extended Data Figure 4.**
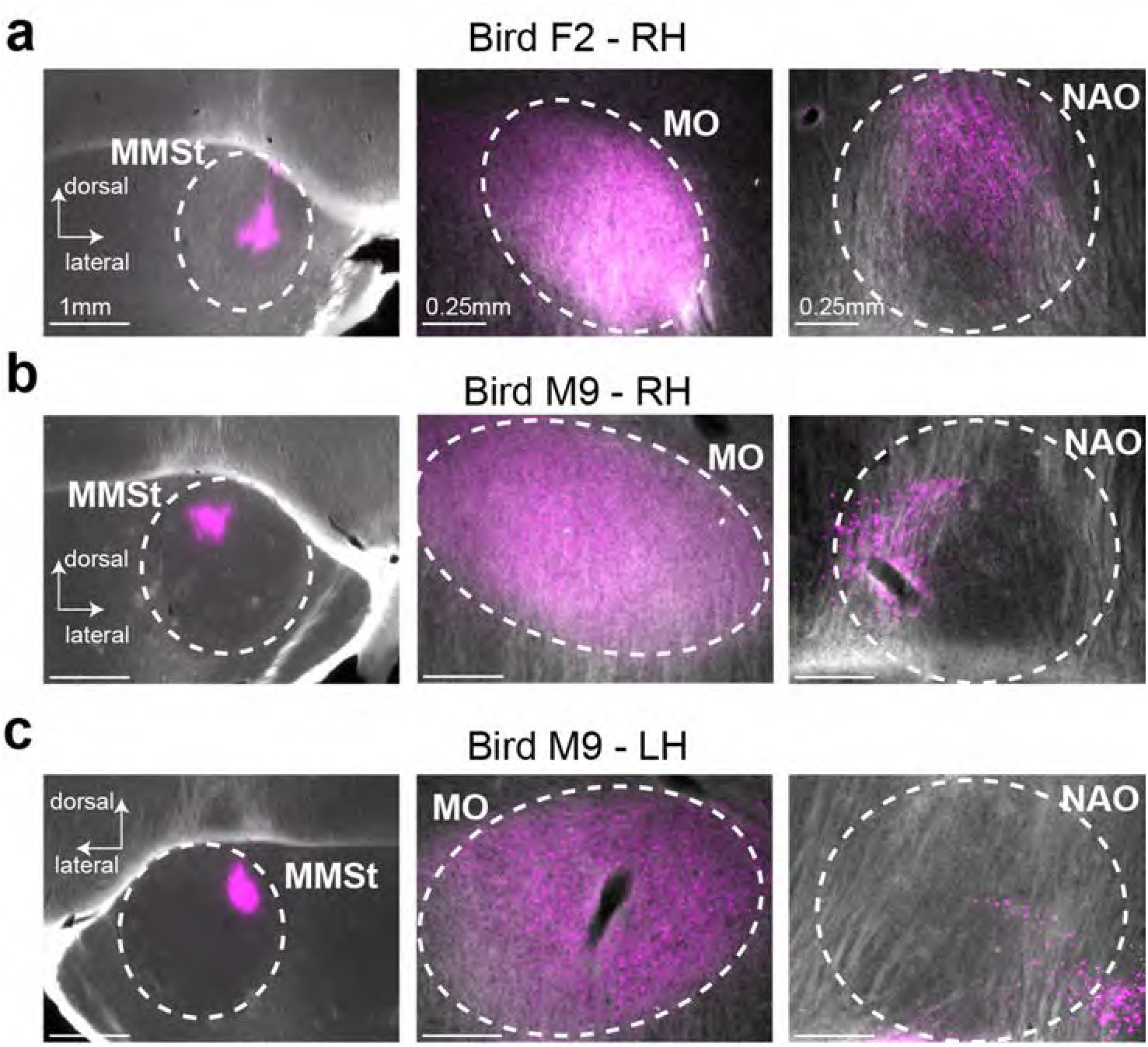
Additional results from the anatomical tracing experiments. a, Focal injection of retrograde tracers in a female budgerigar MMSt (left) backlabeled neurons throughout MO (middle), but only in restricted regions in NAO (right). b, Same as (a), but in the right hemisphere of a male budgerigar. c, Same as in (b), but in the left hemisphere of the same male budgerigar. Images in the same column have the same scale bar: left, 1 mm; middle & right, 0.25 mm. Abbreviations for brain regions from Fig. 3a: AAC, central nucleus of the anterior ar copallium; NLC, central nucleus of the lateral nidopallium; MO, oval nucleus of the anterior mesopallium; NAO, oval nucleus of the anterior nidopallium; MMSt, magnocellular nucleus of the medial striatum; DMM, magnocellular nucleus of the dorsal thalamus; H, hyperpallium; M, mesopallium; N, nidopallium; A, arcopallium; RA, robust nucle us of the arcopallium; HVC, proper name; LMAN, lateral magnocellular nucleus of the anterior nidopallium; MMAN, medial magnocellular nucleus of the anterior nido-pallium; DLM, medial portion of the dorsolateral thalamus; VTA, ventral tegmental area.

**Extended Data Figure 5.**
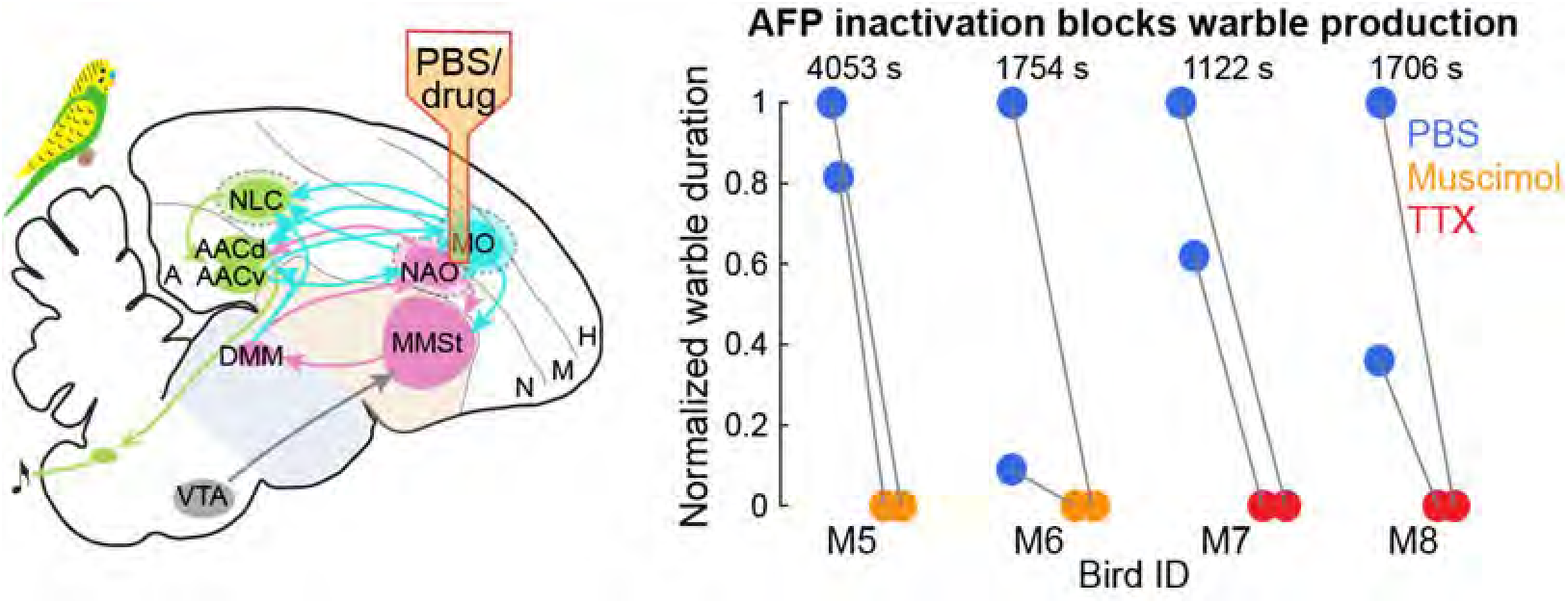
Inactivation of the anterior forebrain pathway (AFP) abolished warble production. Left, schematics of the inactivation experiments using reverse microdialysis. Right, total duration of warble songs produced under the control (blue) or drug (orange/red) condition, normalized to the highest amount of warble produced for a given bird (shown in seconds). Each dot indicates an infusion replicate, with lines connecting the control and drug days that were adjacent.

**Extended Data Figure 6.**
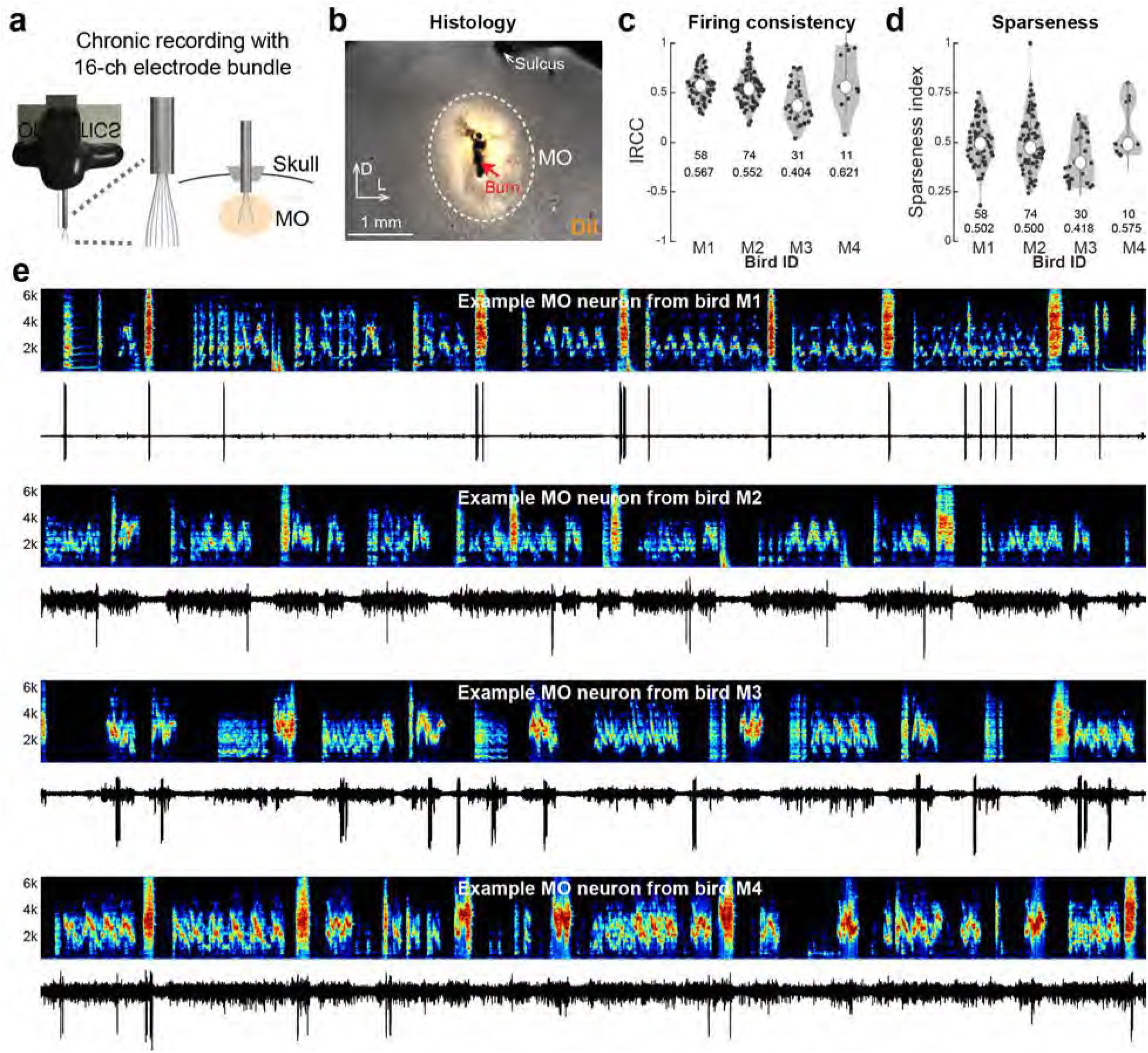
Electrophysiology, histology, and supplementary neural analyses. a, Diagram of the setup for chronic electrophysiological recording in MO. b, Representative histology confirming recording locations in MO. Electrode tracks were labeled with fluorescent dyes before implantation, and recording sites were marked with small electrical lesions after completion of recordings. MO was identified using the anterior sulcus as an anatomical landmark. c, Firing consistency of sparse MO neurons during the production of warble calls, as measured by inter-rendition correlation coefficients (IRCC, Methods) and shown as violin plots. Each black dot indicates one neuron. Large white dots indicate the median value. The first number below the violin indicates the total number of neurons in the analysis, while the second number indicates the mean value. d, Same as in (c), but for the sparseness index (Methods). e, Spectrograms and voltage traces for additional example sparse MO neurons from each bird. Total duration is 10 s for each segment.

**Extended Data Figure 7.**
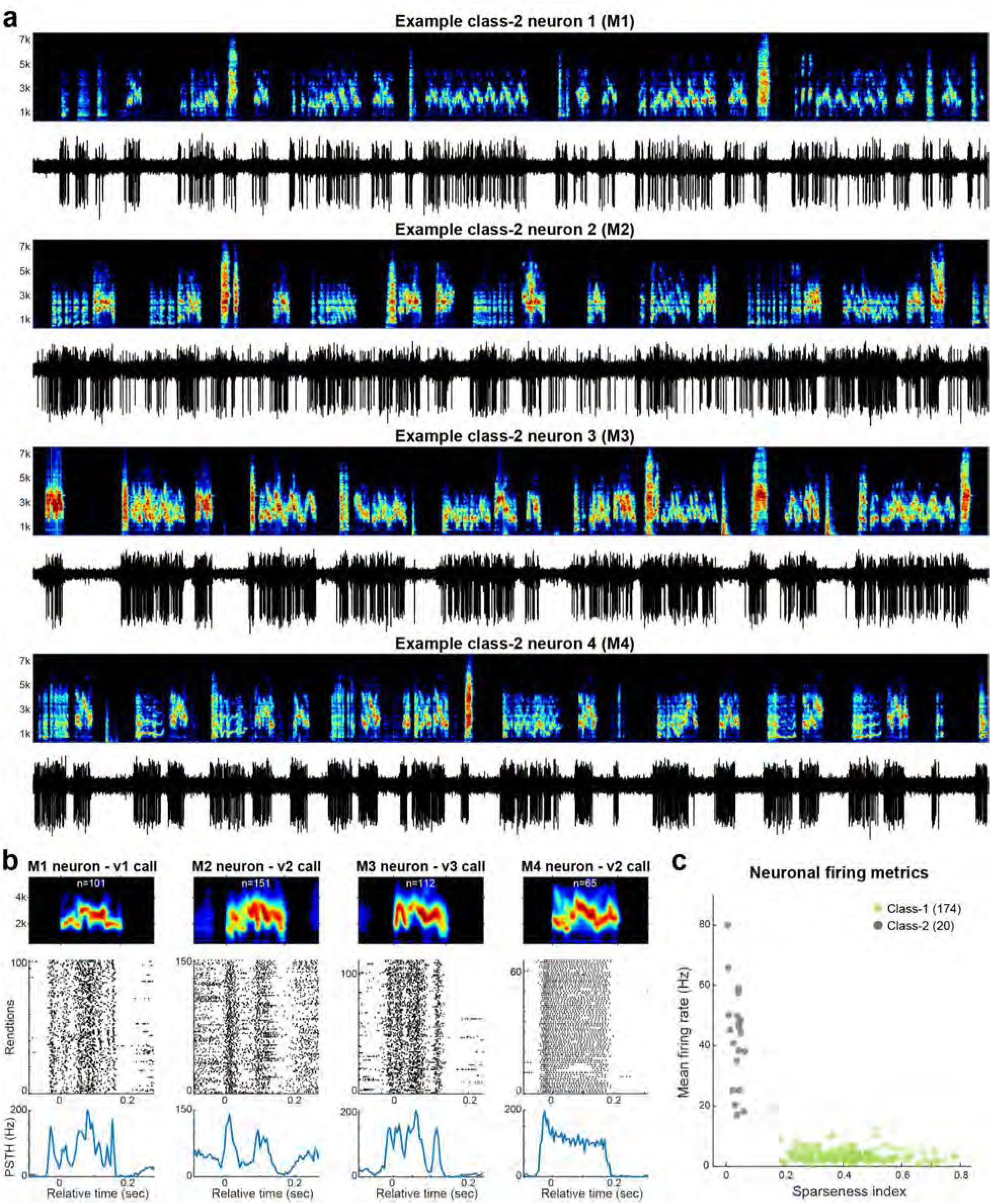
MO neurons that fire broadly. a, Example spectrograms and voltage traces for broad MO neurons from each bird. Total duration is 10 s for each segment. b, Middle, raster plots for the four broad MO neurons shown in (a). Each tick indicates one spike. Spikes from each rendition were aligned to the syllable spike and time-warpped to the mean syllable duration. Top, averaged spectrograms generated from all renditions in the raster plot. Bottom, peristimulus time histogram (PSTH) for spikes shown in the raster plot. c, The sparse (green) and broad (black) MO neurons can be reliably separated by the sparseness index and mean firing rate. Each dot indicates one neuron, colored by the neuronal class.

**Extended Data Figure 8.**
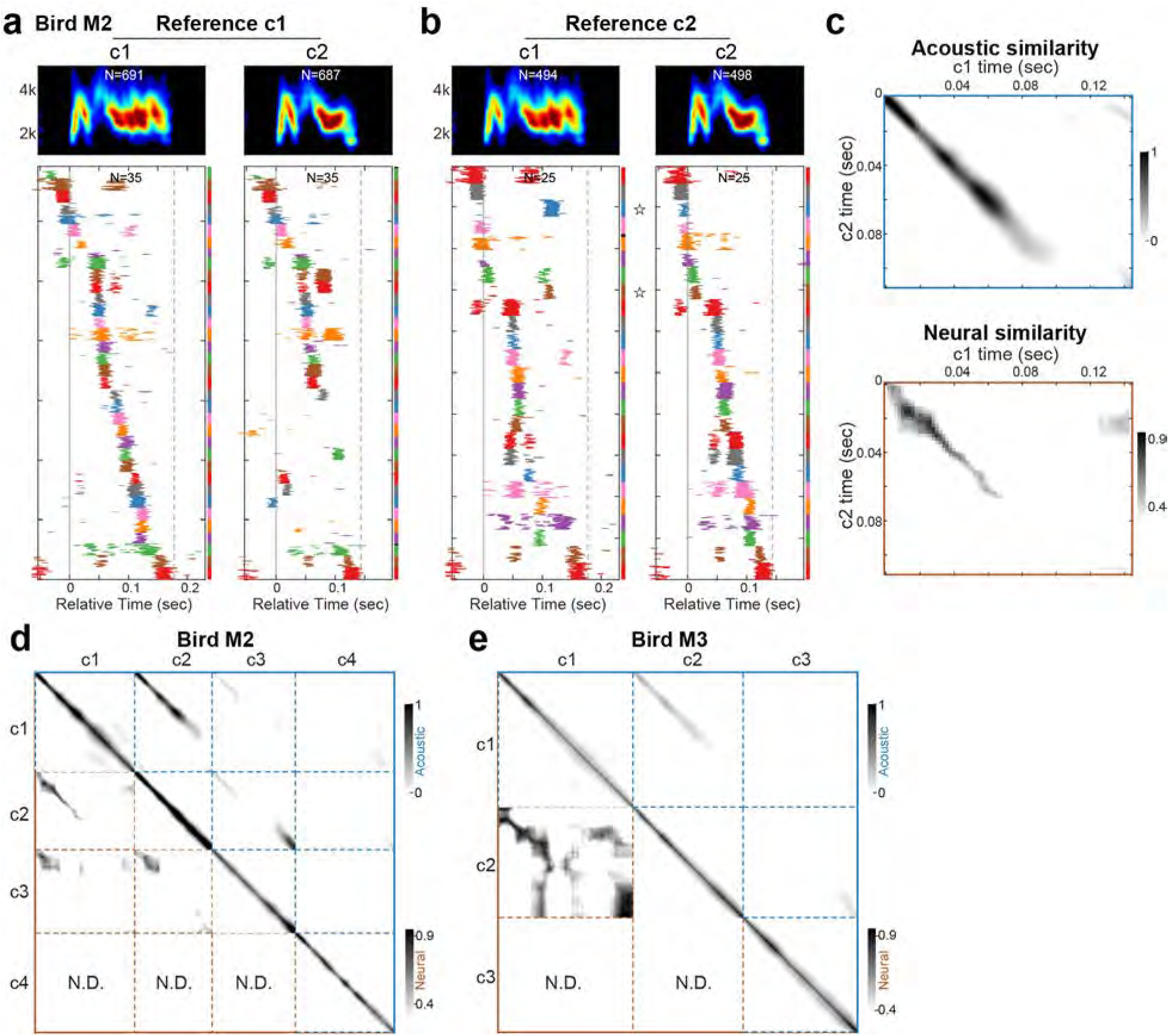
Supplementary analysis for sparse MO neurons that fire during warble call production. a-b, Bottom, population raster plots for sparse MO neurons that burst during the production of warble calls in bird M2, using different call subtypes as the alignment reference. Each row indicates one rendition, with neurons colored differently using 8 colors that are reused periodically. In a given figure panel, the same color on the same row location indicates the same neuron. The colored bars on the right indicate the range of the renditions for a given neuron, with black meaning inadequate rendition numbers. Top, mean spectrograms of all call renditions that were used in the population raster plots below. Numbers indicate the total number of renditions. c, Top, acoustic similarity index between any two time points in call subtype c1 and c2, as measured by dMMD (Methods) and plotted as a matrix. Bottom, neural similarity index between any two time points in call subtype c1 and c2, as measured by SVM decoding accuracy (Methods) and plotted as a matrix. d, The upper right half of the matrix (blue) shows the acoustic similarity, while the lower left half (brown) shows the neural similarity, for all pairwise comparison between call subtypes in bird M2. The neural similarity could be not determined (N.D.) due to the inadequate number of neurons for building a SVM decoding model (Methods). e, Same as in (g), but for bird M3.

**Extended Data Figure 9.**
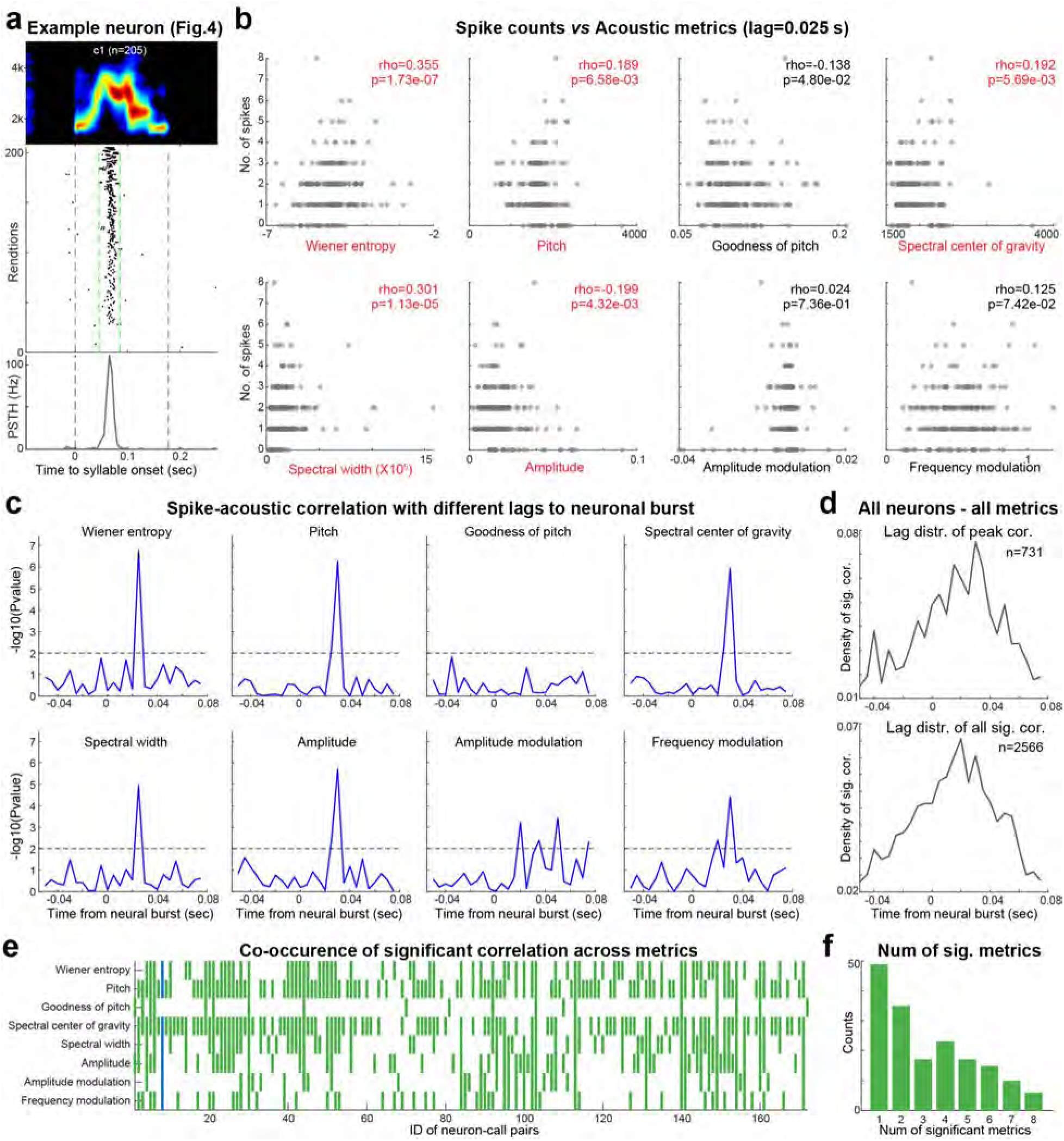
Correlation analysis between spike counts in MO bursts and conventional acoustic metrics. a, Raster plot of an example neuron from bird M1 (same neuron in Figure 4) during the production of warble call subtype c1. Middle, each row indicates one rendition, time-warped to the mean duration of all renditions. Black ticks indicate spikes, gray lines indicate syllable onsets and offsets, green lines indicate the range where spikes were considered within the sparse burst. Top, averaged spectrogram of all renditions in the middle plot. Bottom, peristimulus time histogram (PSTH). b, Scatter plots of spike counts and eight conventional acoustic metrics for the example neuron, at a time lag of 25 ms between the neuronal burst center and a 5 ms sliding window where acoustic metrics were averaged. Each dot indicates one rendition of the call subtype c1. Spearman correlation coefficients and p-values were shown, with red color indicating those with p-values smaller than 0.01. c, Correlation significance between spike counts in the sparse MO bursts and eight acoustic metrics at different lags from −50 ms to +75 ms. P-values from correlation analyses as shown in panel (b) were transformed into negative log scale. Dashed lines indicate the 0.01 significance level. Note that for the example neuron, spike counts are significantly correlated with multiple acoustic metrics, at a positive lag between the neuronal burst and the acoustic sliding window, indicating a pre-motor function. d, Aggregated results of all MO sparse neurons from all birds for all eight acoustic metrics. Across 4 birds, we conducted correlation analysis on all identified call subtypes and MO neurons that exhibited sparse burst (n=299 pairs). For each neuron-call pair, we calculated Spearman correlation between the spike counts in a burst and 8 conventional acoustic metrics with different time lags, which generated 299*8=2392 curves of p-values versus time lags (as shown in panel c). For each curve, we identified all correlations that are significant (p<0.001) and plotted the distribution of their time lags (bottom panel, total 2566 significant points). For curves that have at least one significant correlation, we further identified the peak correlation and plotted the distribution of time lags for these peak correlations (top panel, total 731 peaks). Note that both lag distributions show strong pre-motor bias (lag>0). e, A substantial fraction of neuron-call pairs exhibited significant correlations for multiple acoustic metrics. Each row is one acoustic metric, each column is one neuron-call pair. Green bars indicate significant correlations (p<0.001) within the premotor window (lag=0-50 ms). Blue bars represent results for the example neuron in panel a-c. f, Distribution of number of metrics that correlate significantly with spike counts, for given neuron-call subtype pairs.

**Extended Data Figure 10.**
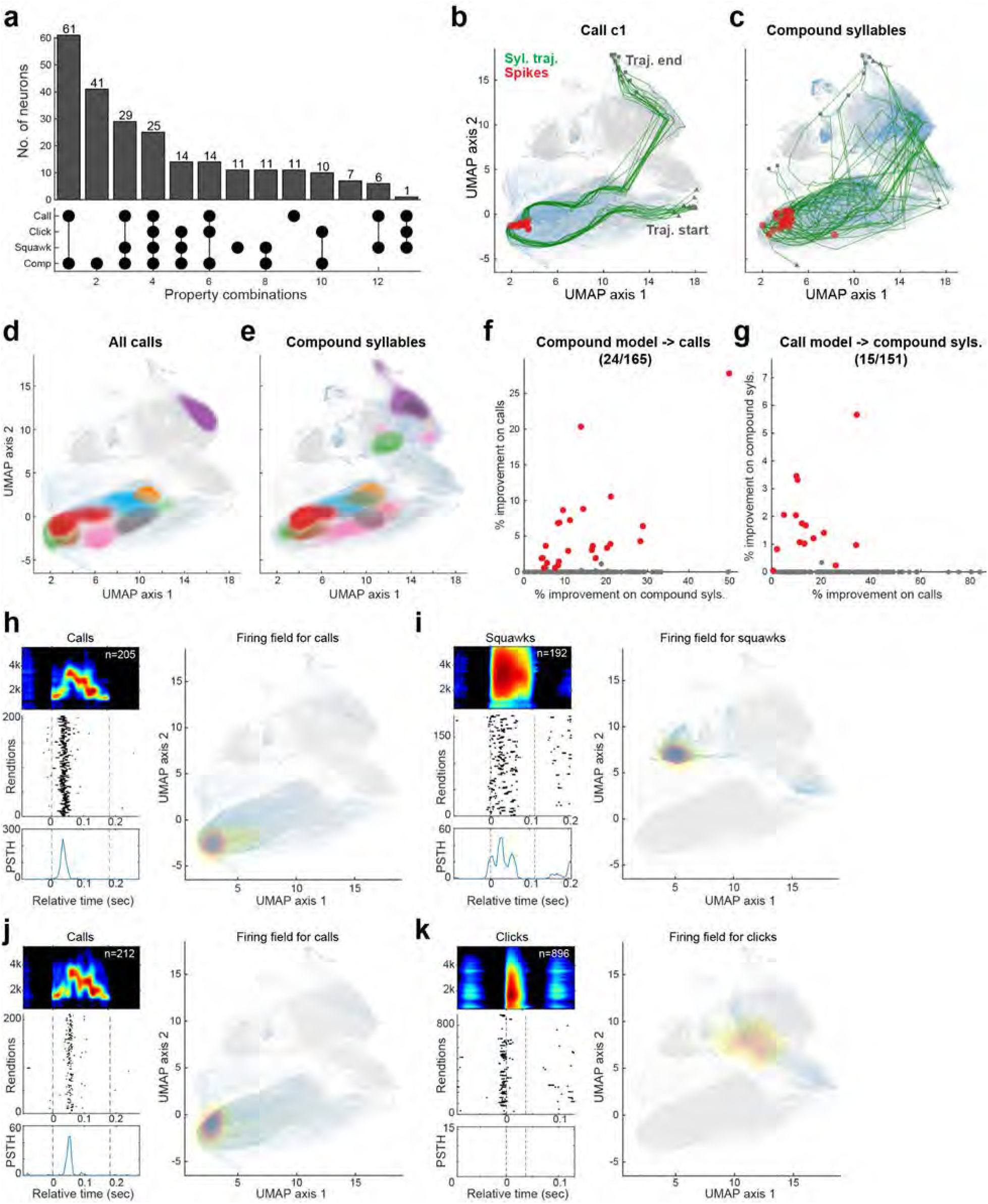
MO neuronal discharge across syllable categories. a, Counts of MO neurons that discharge across syllable types. Top, barplots of MO neuron counts for each firing group. Bottom, neurons were divided into different groups according to which syllable type combinations the neuron exhibited firing (black dots). b-c, Overlay spikes (red dots) of one example MO neuron on the UMAP for call subtype c1 (b) or for compound syllables (c). Green lines indicate syllable trajectories, with grey triangles and squares indicating trajectory starts and ends, respectively. The grey background indicates the embedding of all syllables, while the blue background highlights the embedding of calls (b) or compound syllables (c). d-e, Firing fields of 8 MO neurons during the production of calls (d) or compound syllables (e). Each colored field indicates the density of all spikes for a given neuron. The red neuron is the example neuron in (b-c). Note that the firing fields for these neurons are largely consistent across calls and compound syllables. f, Performance of ensemble learning models that were trained on compound syllables (x-axis) and used to predict firing on calls (y-axis). Each dot indicates one neuron, with red dots indicating neurons whose performance on calls were significantly better than null models (p<0.05, Wilcoxon signed rank test). g, Same as in (f), but for models that were trained on calls (x-axis) and used to predict firing on compound syllables (y-axis). h, One example MO neuron that exhibited discharge during the production of calls and squawks. Left, raster and PSTH, aligned to call onsets. Right, firing field in the UMAP space. Spike density is indicated by the yellow-red color gradient. The grey background indicates the embedding of all syllables, while the blue background highlights the embedding of calls (left) or clicks (right). i, Same neuron as in (h), but for the production of squawks. j-k, Same as in (h-i), but for a different MO neuron that exhibited discharges during the production of calls and clicks.

